# Targeted genomic sequencing of avian influenza viruses in wetlands sediment from wild bird habitats

**DOI:** 10.1101/2023.03.30.534984

**Authors:** Kevin S. Kuchinski, Michelle Coombe, Sarah C. Mansour, Gabrielle Angelo P. Cortez, Marzieh Kalhor, Chelsea G. Himsworth, Natalie A. Prystajecky

## Abstract

Diverse influenza A viruses (IAVs) circulate in wild birds, including dangerous strains that infect poultry and humans. Consequently, surveillance of IAVs in wild birds is a cornerstone of outbreak prevention and pandemic preparedness. Surveillance is traditionally done by testing birds, but dangerous IAVs are rarely detected before outbreaks begin. Testing environmental specimens from wild bird habitats has been proposed as an alternative. These specimens are thought to contain diverse IAVs deposited by broad range of avian hosts, including species that are not typically sampled by surveillance programs. We developed a targeted genomic sequencing method for recovering IAV genome fragments from these challenging environmental specimens, including purpose-built bioinformatic analysis tools for counting, subtyping, and characterizing each distinct fragment recovered. We demonstrated our method on 90 sediment specimens from wetlands around Vancouver, Canada. We recovered 2,312 IAV genome fragments originating from all 8 IAV genome segments. 11 haemagglutinin (HA) subtypes and 9 neuraminidase subtypes were detected, including H5, the current global surveillance priority. Recovered fragments originated predominantly from IAV lineages that circulate in North American resident wild birds. Our results demonstrate that targeted genomic sequencing of environmental specimens from wild bird habitats can be a valuable complement to avian influenza surveillance programs.

## INTRODUCTION

Avian-origin influenza A viruses (AIVs) pose a perennial threat to poultry and human health. Outbreaks in poultry flocks incur significant economic losses^1,2^. They also expose agricultural workers to novel influenza infections, threatening epidemics and global influenza pandemics^3,4,5^. These outbreaks occur when farmlands become contaminated with excreta from infected wild birds. Numerous wild bird species are naturally infected with diverse AIVs, particularly shore birds and waterfowl^6,7^. These birds live in complex communities, resulting in frequent spillovers between species, reassortment of viral genome segments, and emergence of new strains^8,9,10^. Seasonal migrations along intercontinental flyways allow global dissemination of AIVs^11,12^.

Surveillance of AIVs in wild birds is a cornerstone of outbreak prevention and pandemic preparedness^13,14,15,16^. Testing is conducted on live-captured birds, hunter-killed birds, and natural deaths recovered from avian habitats. The objective of these surveillance programs is early detection of strains that are pathogenic to poultry and humans. This would allow agricultural producers to increase biosecurity measures and prevent exposure of livestock to infectious excreta. Due to logistical limitations on the number of birds that can be tested, low detection rates, and sampling biases towards certain avian species, these surveillance programs rarely succeed in forewarning the arrival of dangerous AIVs before outbreaks begin in poultry and humans^15^.

Alternative AIV surveillance strategies have been proposed wherein environmental specimens from wild bird habitats are tested instead of animals^17,18^. The rationale is that AIVs from diverse members of the wild bird community will accumulate in the environment, including AIVs from avian species that are not commonly tested by surveillance programs. Additionally, environmental specimens are comparatively easy to collect and less disruptive to wildlife. Wetlands sediment is one type of environmental specimen in which AIV genomic material has been successfully detected^18,19,20^.

To facilitate AIV surveillance using wetlands sediment, we developed a targeted genomic sequencing method to characterize fragments of influenza A virus (IAV) genome in sediment specimens. The method encompasses three components: 1) a custom panel of hybridization probes targeting all IAV subtypes circulating in avian, swine, and human hosts; 2) sequencing library construction that incorporates a unique molecular index (UMI) on both ends of each genomic fragment in the specimen; and 3) purpose-built bioinformatic tools that resolve UMIs and allow each distinct fragment of IAV genome recovered to be counted and individually characterized.

In this study, we applied our custom method to 90 sediment specimens collected from wetlands around Vancouver, British Columbia, Canada during the autumn and winter of 2021/22. Genome fragments were recovered from varied IAVs, and these fragments were used to assess subtype diversity, host range, and geographic origin of the IAVs in these sediments. Recovered fragments were also used to monitor whether viruses from recognized highly pathogenic AIV (HPAI) clades were present. Further risk assessment was conducted by interrogating recovered fragments for specific genetic markers of virulence.

## RESULTS

### Screening sediment for IAV genomic material by RT-qPCR

Total RNA was extracted from 435 sediment specimens then screened for IAV genomic material by RT-qPCR. 74 sediment specimens (17.0%) were positive. An additional 64 specimens (14.7%) were deemed to be suspect-positive due to having Ct values above the cut-off threshold (n=4) or amplification curves trending towards the critical threshold in the final cycle (n=60). Sequencing capacity was available for 90 specimens. All 74 positive specimens were assayed. 16 randomly chosen suspect-positive specimens were also assayed to assess whether sequencing specimens with indeterminate RT-qPCR results would be worthwhile during future surveillance efforts.

### Detection of IAV genome fragments in sediment by probe capture-based targeted genomic sequencing

IAV genome fragments in these specimens were captured and enriched using a custom panel of hybridization probes. The panel was designed for One Health IAV surveillance, targeting all segments of the IAV genome and providing broadly inclusive coverage of all subtypes circulating in avian, swine, and human hosts (Table S1 and Table S2). Captured material was sequenced (Table S3) then analyzed with two purpose-built bioinformatic tools called HopDropper and FindFlu. HopDropper uses unique molecule index (UMI)-based anaylsis to generate consensus sequences for each distinct fragment of IAV genome recovered^21^. FindFlu characterizes these fragment consensus sequences and determines the IAV genome segments from which they originated.

We detected 2,312 IAV fragments in specimens that were positive by RT-qPCR (Figure 1A). Only 8 IAV fragments were detected in suspect-positive specimens. Low recovery from specimens with indeterminate RT-qPCR results indicated that future surveillance activities should focus on positive specimens only. To reflect surveillance based only on specimens positive by RT-qpCR, the 16 suspect-positive specimens and the 8 IAV genome fragments recovered from them were omitted from the following analyses.

**Figure 1:**
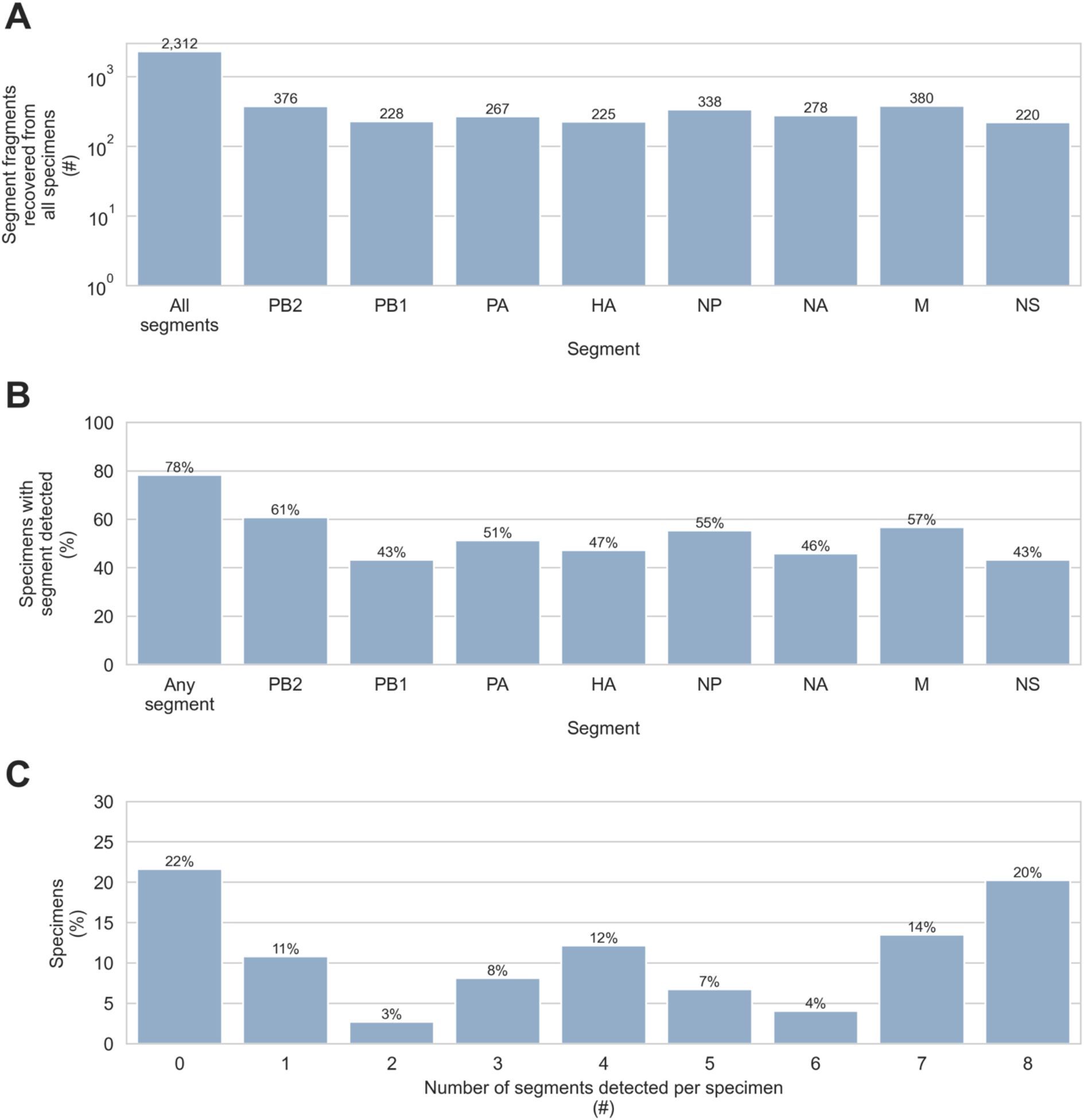
Detection of influenza A virus genome fragments in sediment by probe capture-based targeted genomic sequencing. Influenza A virus (IAV) genome fragments were recovered from 74 sediment specimens that tested positive for IAV genomic material by RT-qPCR. **A)** The number of IAV genome fragments recovered from all specimens was counted. In addition to the total count, the number of fragments originating from each of the 8 IAV genome segments (PB2, PB1, PA, HA, NP, NA, M, and NS) was also determined. **B)** The sensitivity of probe capture-based targeted genomic sequencing was determined for specimens that tested positive by RT-qPCR. Overall sensitivity was calculated as the percentage of specimens positive by RT-qPCR where probe capture-based targeted genomic sequencing detected at least one IAV genome fragment from any genome segment. Sensitivity was also calculated for each of the IAV genome segments separately. **C)** The number of different IAV genome segments detected in each specimen was determined.

IAV genomic material was detected by probe capture-based sequencing in 58 of 74 specimens (78%) that tested positive by RT-qPCR, and fragments from all 8 IAV genome segments were recovered (Figure 1AB). IAV fragments were not evenly distributed across specimens, however. The three specimens with the most IAV fragments contained 809, 246, and 148 IAV fragments respectively. Collectively these three specimens yielded 56% of all IAV fragments detected. The median specimen contained only 6 IAV fragments (Figure 2A). Furthermore, only 20% of specimens contained fragments from all 8 genome segments (Figure 1C), and no individual genome segment was detected in more than 61% of specimens (Figure 1B).

**Figure 2:**
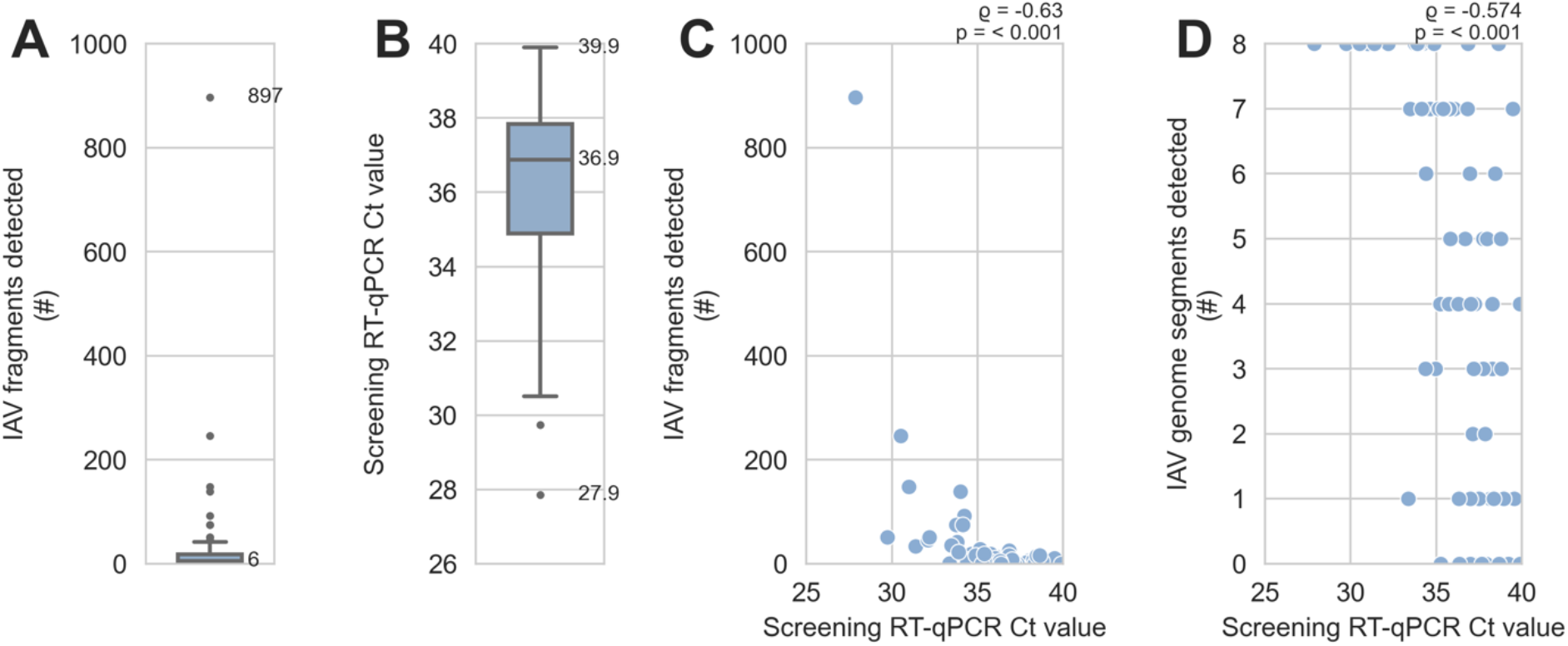
Detection of influenza A virus genome fragments was limited by low abundance of viral genomic material in sediment specimens. 2,312 fragments of influenza A virus (IAV) genome were recovered from 74 sediment specimens that tested positive for IAV genomic material by RT-qPCR. **A)** The number of IAV genome fragments recovered per specimen was counted. This distribution includes specimens where no IAV fragments were recovered. The median and maximum are indicated. **B)** Distribution of screening RT-qPCR Ct values for specimens, including specimens where no IAV fragments were recovered. The minimum, median, and maximum are indicated. **C)** There was a moderate and statistically significant monotonic association between screening RT-qPCR Ct values and the number of IAV genome fragments detected by probe capture-based targeted genomic sequencing. Results of Spearman’s rank correlation are indicated above the upper-right corner of the scatterplot. **D)** There was a moderate and statistically significant monotonic association between screening RT-qPCR Ct values and the number of different IAV genome segments detected by probe capture-based targeted genomic sequencing. Results of Spearman’s rank correlation are indicated above the upper-right corner of the scatterplot.

These results, together with the high Ct values observed when screening specimens by RT-qPCR (Figure 2B), suggested that low abundance of viral material in these specimens caused stochastic, incomplete recovery by probe capture. Indeed, there was a statistically significant monotonic association between lower Ct values (*i.e.* greater abundance of viral genomic material) and higher numbers of IAV fragments detected (Figure 2C). Lower Ct values were also significantly associated with the detection of more of the IAV genome segments (Figure 2D).

### Diversity of IAV subtypes detected

Subtyping the haemagglutinin (HA) and neuraminidase (NA) genome segments is central to IAV surveillance and diagnosis, so our bioinformatic tool FindFlu automatically reports the subtypes of all HA and NA fragments identified. We observed a high diversity of HA and NA subtypes in the 74 sediment specimens that tested positive for IAV material by RT-qPCR. 11 of the 16 avian-origin HA subtypes were detected, and all 9 of the avian-origin NA subtypes were detected (Figure 3AB). The most widespread HA and NA subtypes were H6 and N2. These were present in 15% and 26% of specimens respectively (Figure 3A). The most abundant HA and NA subtypes in terms of total fragments detected were H5 and N2. The number of fragments detected for these subtypes were 93 and 130 respectively (Figure 3B).

**Figure 3:**
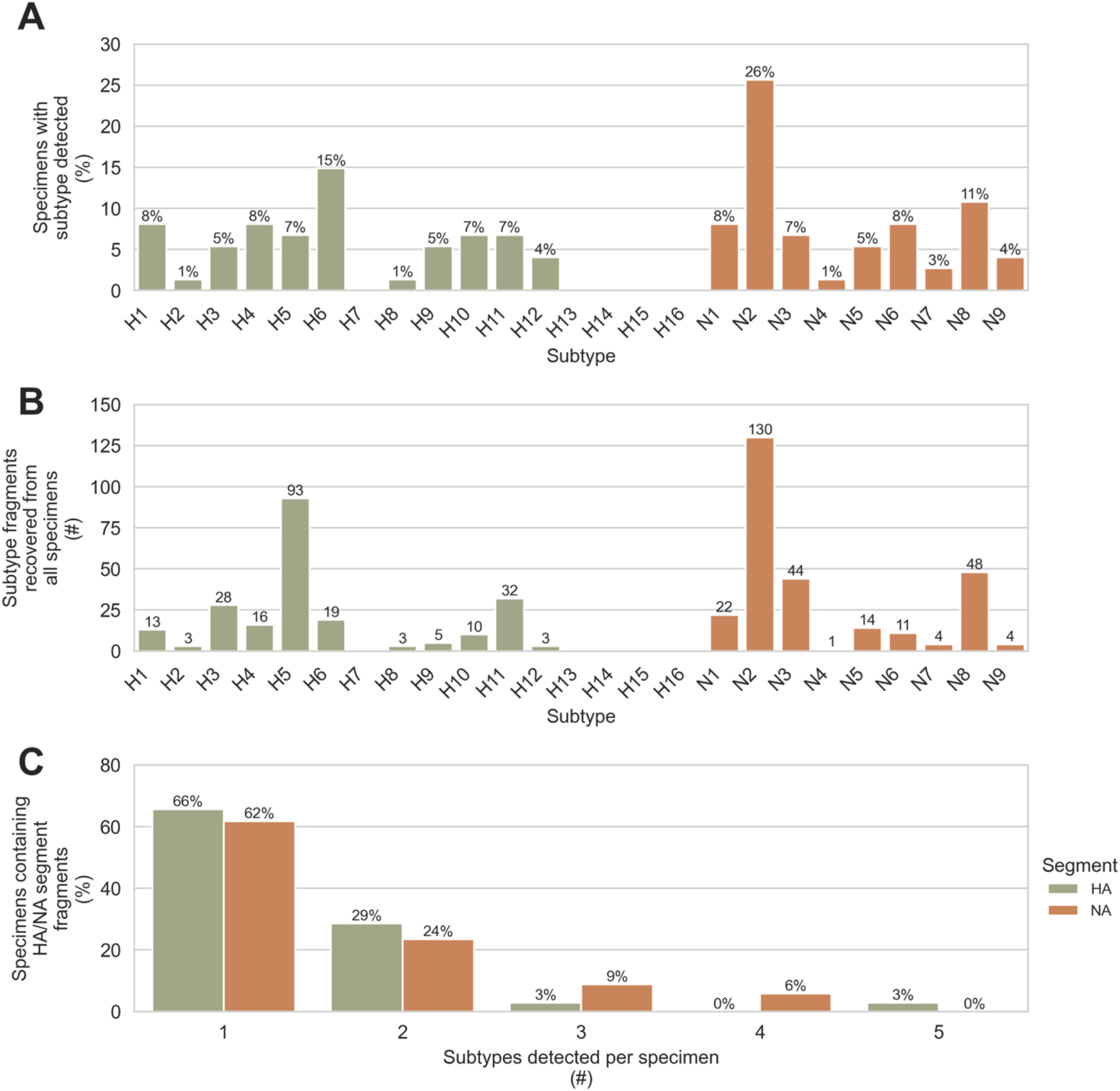
Diverse haemagglutinin and neuraminidase subtypes were detected in wetlands sediment using probe capture-based targeted genomic sequencing. Fragments of the haemagglutinin (HA) and neuraminidase (NA) genome segments were recovered from 74 sediment specimens that tested positive for influenza A virus (IAV) genomic material by RT-qPCR. 225 HA fragments were recovered from 35 specimens, and 278 NA fragments were recovered from 34 specimens. **A)** The percentage of specimens containing each HA and NA subtype was determined. **B)** The total number of HA and NA fragments recovered for each HA and NA subtype was counted. **C)** The number of different HA subtypes detected in each HA-positive specimen was determined, and the number of different NA subtypes detected in each NA-positive specimen was determined.

One of the proposed advantages of using environmental specimens for surveillance is the possibility that individual specimens might contain diverse viruses deposited by multiple hosts. We assessed this by counting the number of different HA or NA subtypes present in the same specimen (Figure 3C). Up to 5 different HA or NA subtypes were observed in the same specimen. 34% of HA-positive specimens contained more than one HA subtype, and 38% of NA-positive specimens contained more than one NA subtype.

### Assessing confidence in detections based on limited numbers of recovered genome fragments

Many of the segment/subtype detections in this study were based on the presence of a limited number of fragments (Figure 2A and Figure S1). This suggested the possibility of false detections. First, we considered whether some of these detections were caused by demultiplexing artefacts, *e.g.* mutations or base calling errors in library barcodes that occasionally caused limited numbers of IAV reads to be misassigned to incorrect libraries. To assess this, we determined the number of read pairs that described each IAV fragment (Figure 4). The median number of read pairs per IAV fragment was 852, and 90% of all IAV fragments were described by at least 167 read pairs. Based on these high read pair counts, the actual presence of these IAV fragments in their assigned libraries was strongly supported.

**Figure 4:**
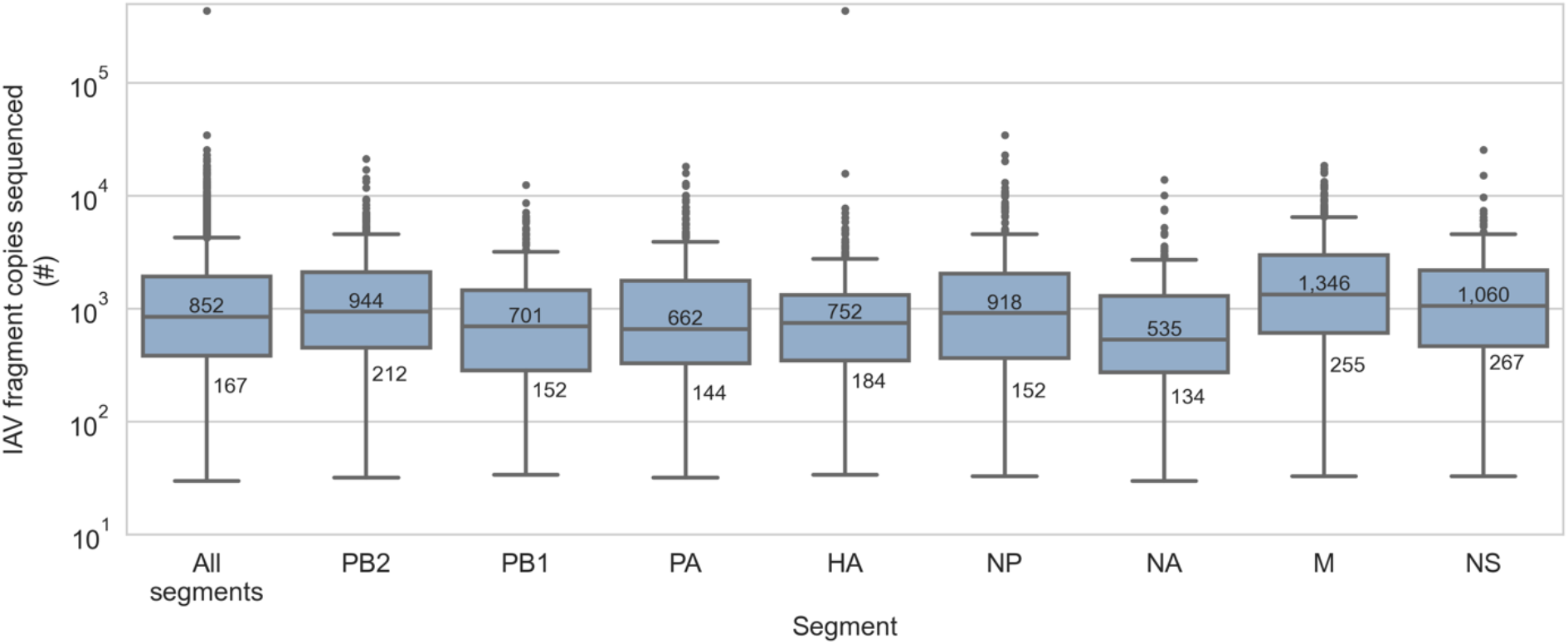
Recovered fragments of influenza A virus genome were sequenced deeply. Influenza A virus (IAV) genome fragments were recovered from 74 sediment specimens that tested positive for IAV genomic material by RT-qPCR. Multiple copies of each IAV fragment were sequenced, increasing sequencing depth per fragment. The median and 10th percentile of copies sequenced per fragment are indicated overall and for each IAV genome segment.

Next, we considered cross-contamination of IAV genomic material between specimens during laboratory handling as a source of false detections. When designing this custom targeted genomic sequencing method, we anticipated the potential for cross-contamination between specimens and incorporated strategies to mitigate this risk. First, the positive control target for this method is a synthetic oligomer with an artificial, computer-generated sequence that does not resemble IAV or any other organism. This ensures that positive controls do not contaminate surveillance specimens with exogenous IAV genomic material. Second, negative controls are composed of commercially prepared human reference RNA background material. Unlike typical water blanks, these contain sufficient total RNA mass for robust library construction, thereby providing more sensitive detection of low-level cross-contamination. No IAV fragments were observed in any of the 6 negative controls processed alongside sediment specimens in this study (Table S3). Taken together, this method design and these control specimen results indicated that cross-contamination was not a measurable source of false detections in this study.

Finally, we considered whether index hopping had attributed detections to incorrect libraries. Index hopping is a form of chimeric PCR artefact where library molecules acquire the barcodes of another library during pooled amplification reactions^22,23,24^. We anticipated index hopping during the post-capture PCR step of this method for three reasons. First, libraries are pooled for capture, so a variety of library barcodes are present on template molecules in the post-capture PCR. Second, the low abundance of viral genomic material in these libraries requires extensive amplification during the post-capture PCR. Third, the post-capture PCR provides favourable conditions for chimera formation because of the numerous amplification cycles, low abundance and complexity of captured material, and fragmented condition of viral genomes.

To identify index hops and other chimeric artefacts, we adopted library construction techniques that associate a unique molecular index (UMI) with both ends of each genomic fragment. This was combined with paired end sequencing on captured material to identify the pair of UMIs associated with each sequenced molecule. A purpose-built bioinformatic tool called HopDropper, which analyzes the frequency and co-occurrences of UMIs, was used to discard sequencing reads originating from potential chimeras and index hops. To confirm the removal of chimeras and index hops by HopDropper, we performed two independent probe captures on the pool of libraries prepared from these specimens, then we separately analyzed each capture with HopDropper. We reasoned that UMIs enriched by both captures should de-multiplex to the same library and be paired with the same other UMI after each capture. 2,191 UMIs were enriched in both replicates. 2,172 of these UMIs (99.1%) were de-multiplexed to same library in both replicates, and 2,148 of these UMIs (98.0%) were paired with the same other UMI in both replicates. This indicated that chimeric artefacts formed during post-capture PCR were largely absent following analysis by HopDropper, and that index hopping was not responsible for systematic false IAV detections in this study.

### Length of IAV fragments recovered by probe capture-based targeted genomic sequencing

FindFlu determines the segment and subtype of IAV fragments by aligning them to IAV reference sequences. It also uses these alignments to estimate the length of each recovered IAV fragment. For these specimens, the median IAV fragment length was 370 nucleotides, but lengths ranged from 104 to 2,113 nucleotides (Figure 5A). FindFlu also uses these estimated fragment lengths to calculate how much each recovered fragment covers of its best-matching reference sequences (Figure 5B). In this study, the median IAV fragment represented only 24% of the segment from which it originated, but some fragments represented up to 99% of their segment of origin.

**Figure 5:**
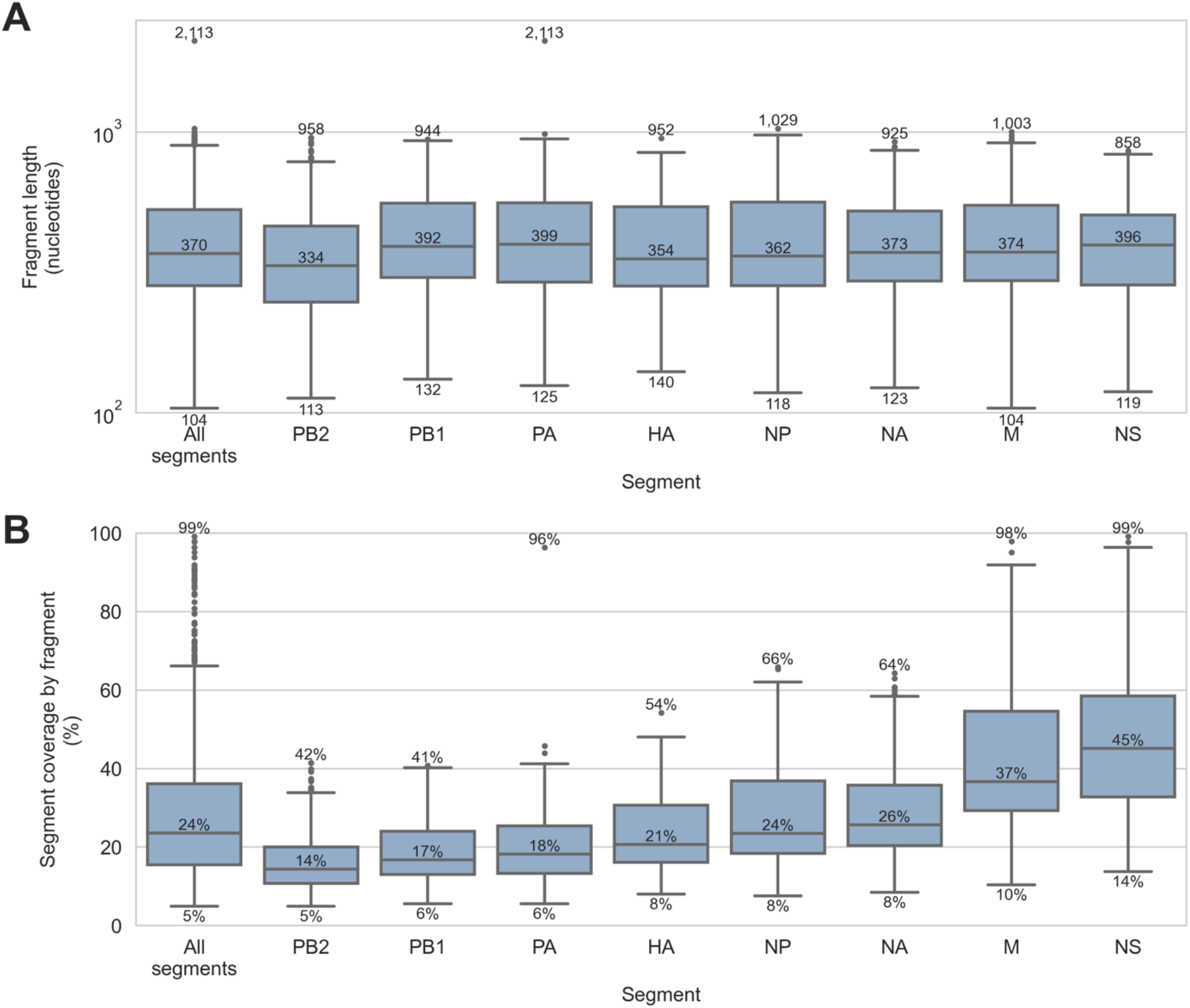
Length of influenza A virus genome fragments recovered from sediment specimens by probe capture-based targeted genomic sequencing. Influenza A virus (IAV) genome fragments were recovered from 74 sediment specimens that tested positive for IAV genomic material by RT-qPCR. **A)** The length of each IAV genome fragment was estimated by FindFlu, a tool that aligned fragment sequences to a database of 555,364 IAV reference sequences (collected globally from avian, swine, and human hosts). Fragment length estimates were calculated from the start and end coordinates of these alignments. **B)** FindFlu also estimated how much each fragment covered of its segment of origin by dividing the estimated fragment length by the length of the reference sequences to which it aligned.

### Characterizing potential host range and geographic origin of IAVs in sediment

When assessing zoonotic risks to agriculture and public health, segment and subtype identification are often insufficient. It is important to know if detected IAVs are similar to those that have previously spilled over into poultry or humans. It is also crucial to identify incursions of viruses from regions where pathogenic strains are known to circulate. For these reasons, the reference sequences used by FindFlu are annotated with the host species from which they were collected and the country where the collection occurred. FindFlu reports these annotations for the best-matching reference sequences of each IAV genome fragment. This provides a qualitative characterization of the potential host range and geographic distribution of the IAVs from which the recovered fragments originated.

In this study, strong alignments were obtained between recovered IAV fragments and FindFlu reference sequences (Figure S2). Median alignment identity was 99.0% and median alignment query coverage was 99.8%. This indicated that the IAVs detected in these sediment specimens were very similar to previously described IAVs, so their characteristics could be confidently inferred from the annotations of their best-matching FindFlu reference sequences.

Recovered fragments had their closest matches to IAVs that were pre-dominantly isolated from North American resident waterfowl and shorebird species (Figure 6 and Figure 7). We noted some fragments were most similar to IAVs observed in Eurasia, *e.g.* H3s from Japan and Mongolia and H6s from South Korea (Figure 7), reflecting intercontinental migration of AIV hosts and the potential for detecting incursions of Eurasian viruses into the Americas. We also noted that small minorities of PB1, PA, N3, and M fragments had their best alignments to reference sequences collected from chickens and turkeys (Figure 6). Furthermore, 5% of PB1 segment fragment best alignments were to a reference sequence collected from humans (Figure 6). This PB1 segment reference sequence (GenBank accession CY125726) was collected from a Mexican poultry worker infected with zoonotic H7N3 IAV in 2012^25^.

**Figure 6:**
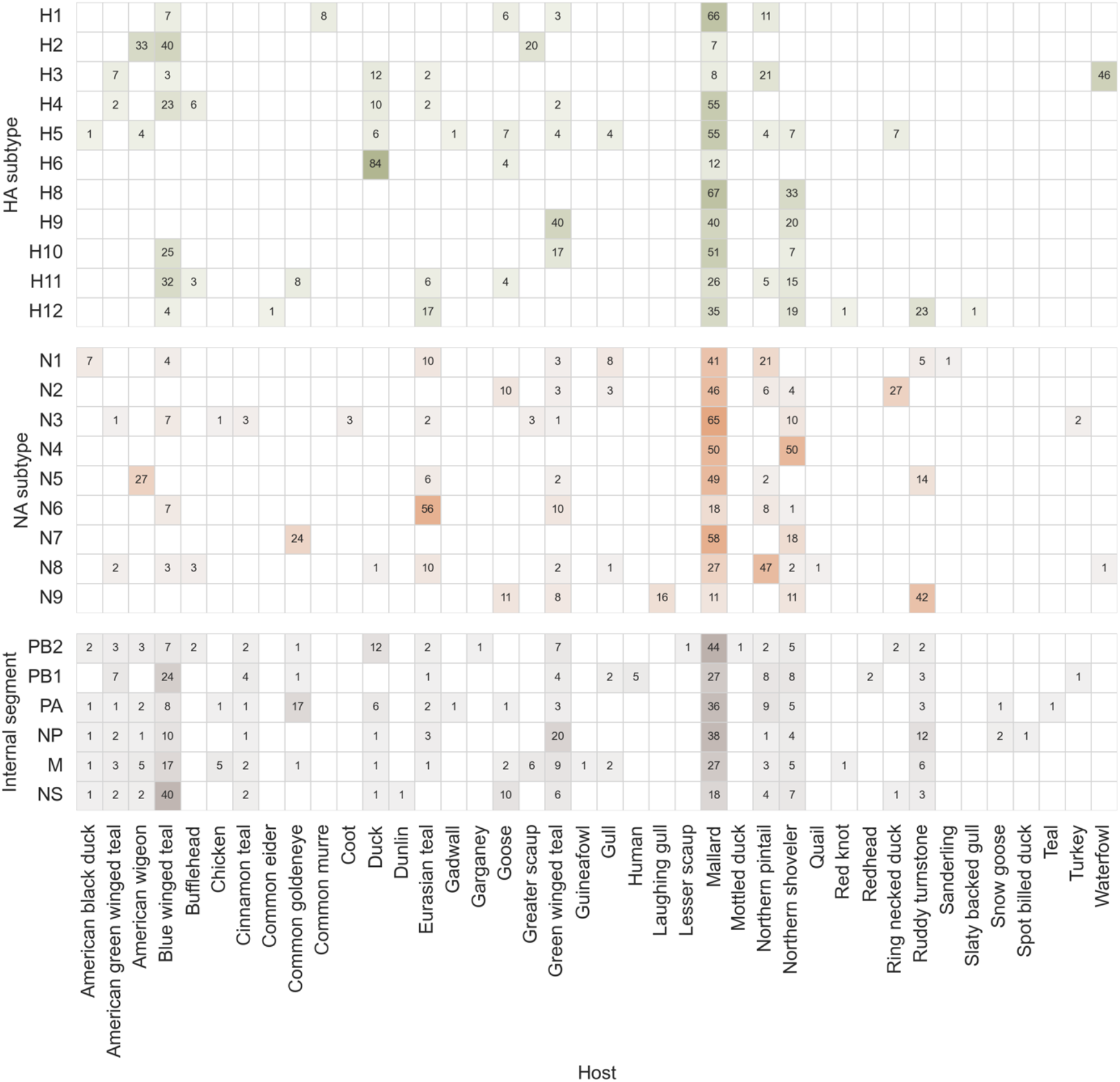
Inferred host species range of influenza A viruses detected in wetlands sediment by probe capture-based targeted genomic sequencing. Influenza A virus (IAV) genome fragments were recovered from 74 sediment specimens that tested positive for IAV genomic material by RT-qPCR. Host species range was inferred for each fragment by FindFlu, a tool that aligned IAV fragment sequences to a database of 555,364 IAV reference sequences (collected globally from avian, swine, and human hosts). Each reference sequence was annotated with the host species from which it was collected. Numbers inside cells indicate the percentage of IAV fragments from a particular segment/subtype whose best-matching reference sequences were collected from the corresponding host species. When fragments had multiple best-matching reference sequences with different host species annotations, fractions of those fragments were proportionally allotted to each host species. Percentages less than 1% were not reported.

**Figure 7:**
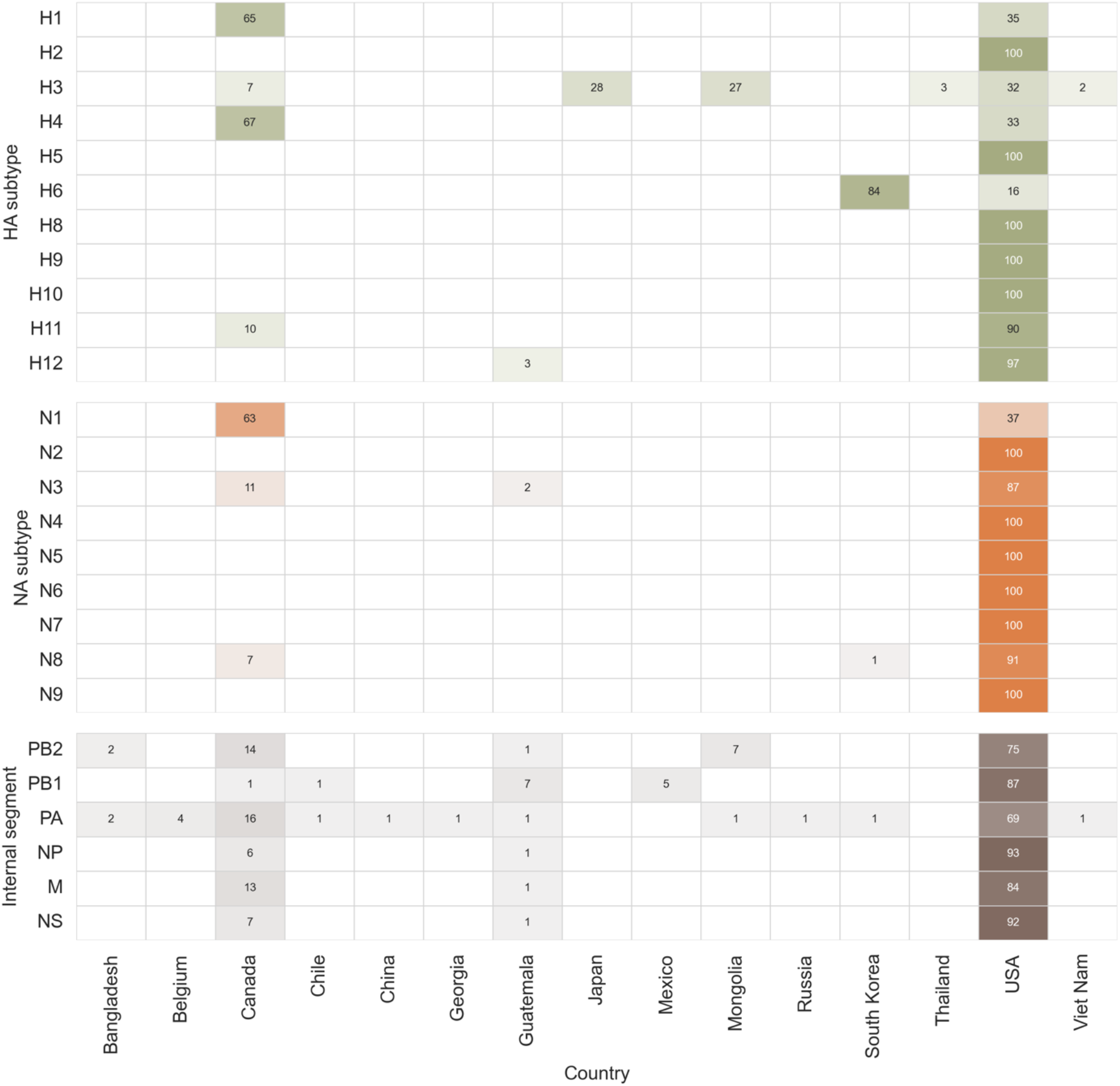
Inferred geographic range of influenza A viruses detected in wetlands sediment by probe capture-based targeted genomic sequencing. Influenza A virus (IAV) genome fragments were recovered from 74 sediment specimens that tested positive for IAV genomic material by RT-qPCR. Geographic range was inferred for each fragment by FindFlu, a tool that aligned IAV fragment sequences to a database of 555,364 IAV reference sequences (collected globally from avian, swine, and human hosts). Each reference sequence was annotated with the country in which it was collected. Numbers inside cells indicate the percentage of IAV fragments from a particular segment/subtype whose best-matching reference sequences were collected in the corresponding country. When fragments had multiple best-matching reference sequences with different country annotations, fractions of those fragments were proportionally allotted to each country. Percentages less than 1% were not reported.

### Assessing the presence of highly pathogenic goose/Guangdong/96 lineage H5 viruses

Next, we focused our analysis on H5 fragments due to the global importance of the highly pathogenic goose/Guangdong/96 (gs/Gd) lineage^26^. Viruses in this H5 lineage have caused numerous outbreaks in poultry and humans since it emerged in the mid-1990s. To identify these threats, our custom One Health IAV probe panel was designed to provide extensive coverage of gs/Gd clades (Table S2), and H5 reference sequences used by FindFlu are further annotated with their H5 lineage and clade. None of the H5 fragments we recovered had their best alignments to gs/Gd reference sequences; all best matches were to viruses belonging to American non-gs/Gd lineages (Figure 8A). When specimen collection for this study began, gs/Gd viruses had not been detected in North America since the end of a previous epizootic in 2015^27^, but they were an escalating problem across Eurasia^28,29,30,31^. No incursions of Eurasian H5s were detected in this study; all recovered H5 fragments had their best alignments to viruses collected in North America (Figure 8B). Finally, none of the recovered H5 fragments had their best alignments to IAVs that were collected from poultry or humans (Figure 8C). All best alignments were to viruses collected from waterfowl and shorebirds.

**Figure 8:**
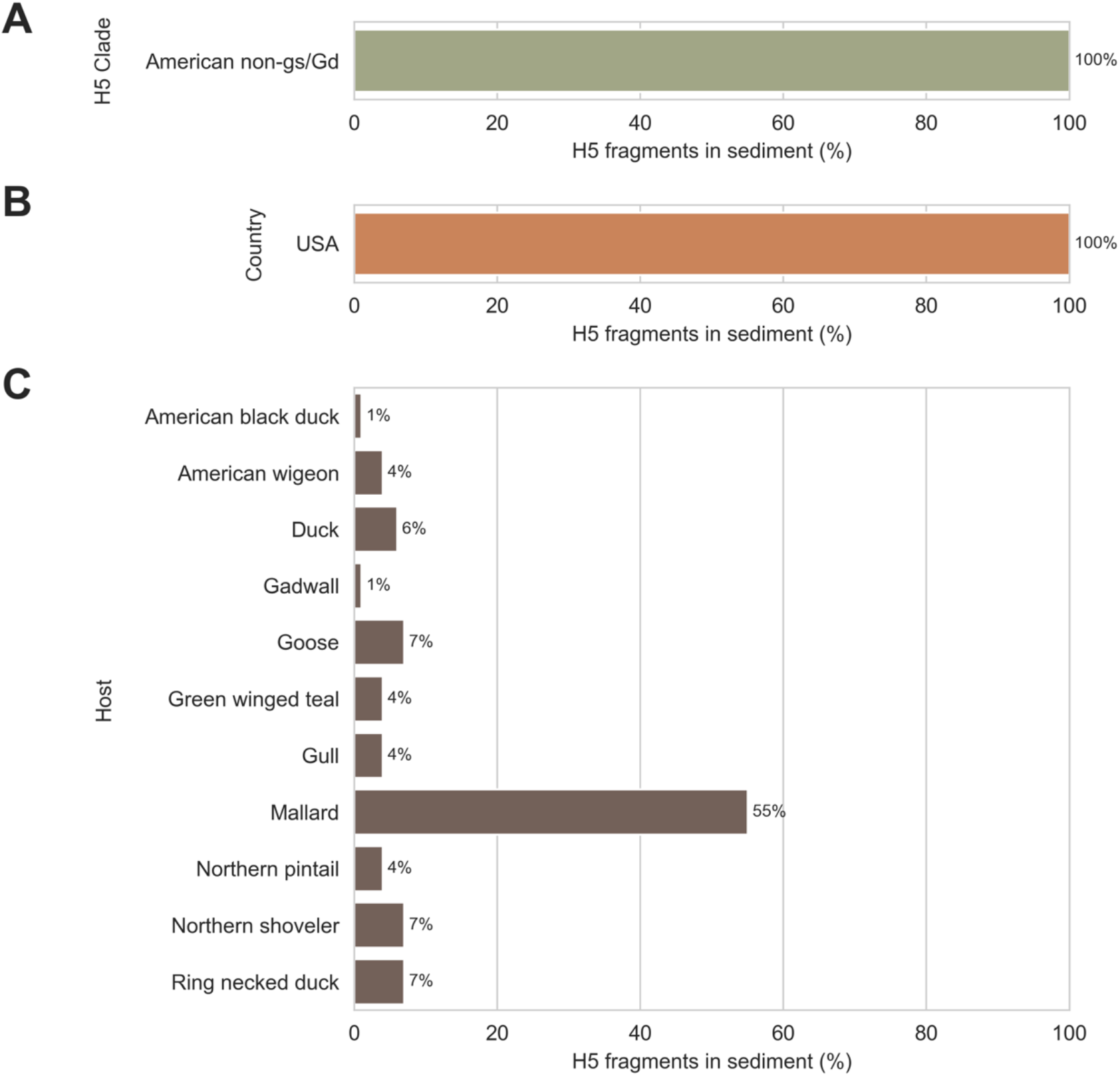
Inferred clade, host species range, and geographic range of H5 subtype influenza A viruses detected in wetlands sediment. 93 fragments of H5 subtype haemagglutinin (HA) genome segment were recovered from 74 specimens that tested positive for influenza A virus (IAV) genomic material by RT-qPCR. Clade, host species range, and geographic range were inferred for each H5 fragment by FindFlu, a tool that aligned IAV fragment sequences to a database of 555,364 IAV reference sequences (collected globally from avian, swine, and human hosts). This database included 6,041 H5 subtype HA segment reference sequences. **A)** All H5 fragments had their best matches to reference sequences belonging to American non-goose/Guangdong (gs/Gd) lineages. **B)** All H5 fragments had their best matches to reference sequences collected in the United States of America (USA). **C)** All H5 fragments had their best matches to reference sequences collected from waterfowl and shorebird species.

We also evaluated the phylogenetic relationship of the H5 viruses in these specimens to the gs/Gd lineage. Direct phylogenetic comparison was complicated by the fragmentary and incomplete sequences recovered from the sediment. We deliberately did not attempt to assemble fragments into larger contigs; since there was evidence of multiple viruses in many of these specimens (Figure 3C), we did not want to inadvertently create chimeric H5 segment sequences.

Instead, we constructed a proxy phylogenetic tree of H5 reference sequences, then we aligned the H5 fragments we recovered to these reference sequences to situate the fragments in their phylogenetic context (Figure 9). This analysis indicated that the H5 IAVs we detected were only distantly related to gs/Gd viruses, diverging from each other before the common ancestor of all gs/Gd lineage IAVs emerged in the mid-1990s.

**Figure 9:**
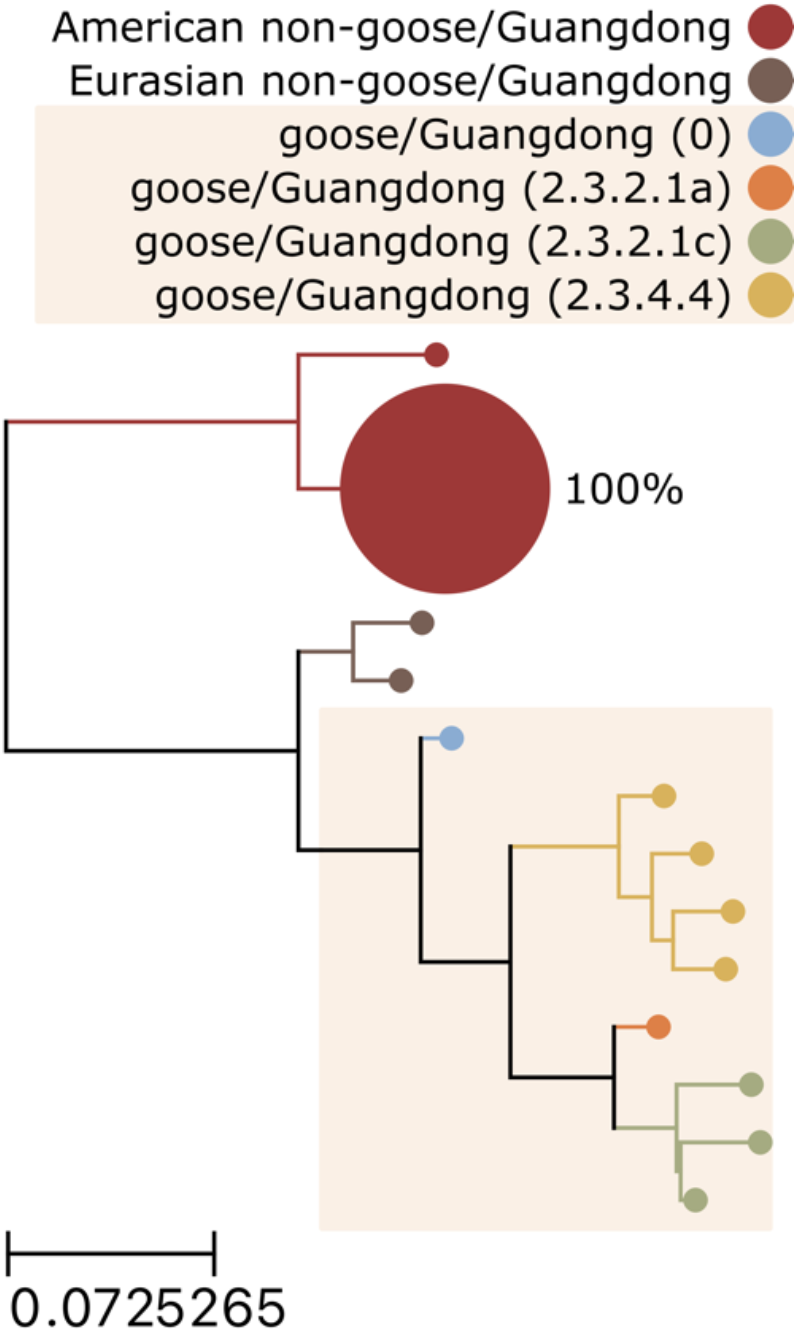
Phylogenetic context of H5 subtype influenza A viruses detected in wetlands sediment by probe capture-based targeted genomic sequencing. A proxy phylogenetic tree was constructed from 147 recent haemagglutinin (HA) segment nucleotide reference sequences belonging to the H5 subtype. Reference sequences were collected globally since 2018 (the past 5 years, inclusive). The HA segment sequence from the prototypical goose/Guangdong/96 lineage (GenBank accession NC_007362) was also included to represent Clade 0 of this lineage. Monophyletic groups of highly similar sequences (all leaves within 0.025 substitutions/site of their common ancestor) were collapsed into single leaves for visual clarity. Leaves were coloured according to their H5 lineage and clade. Background shading was applied to Gs/Gd lineage clades. 93 fragments of H5 subtype HA segments were recovered from sediment specimens. These H5 fragments were aligned to the reference sequences composing the proxy tree. For each tree leaf, the percentage of recovered H5 fragments whose best-matching reference sequences belonged to the leaf was calculated. These percentages were indicated beside each leaf and used to scale leaf sizes.

Finally, we assessed the virulence of the H5 IAVs in the sediment by characterizing HA cleavage sites on recovered fragments. A common feature of HPAI is the presence of polybasic amino acid insertions in this cleavage site^30,32^. We identified 9 H5 fragments on which the HA cleavage site had been sequenced (Table 1). All 9 of these fragments contained the same canonical low-pathogenicity H5 cleavage site motif: PQRETRGLF.

**Table 1:** Haemagglutinin cleavage site on H5 subtype fragments recovered from sediment specimens had canonical low-pathogenicity motifs. 93 fragments of influenza A virus (IAV) genome were recovered that originated from H5 subtype haemagglutinin (HA) genome segments. Fragments were translated and aligned to the HA segment of the prototypical goose/Guangdong/96 lineage strain (GenBank accession NC_007362). HA cleavage site motifs were extracted from translated, aligned sequences on fragments that spanned the cleavage site.

| H5 cleavage site motif | H5 fragments containing motif | Total H5 cleavage site-spanning fragments | Percent of total (%) |
| --- | --- | --- | --- |
| PQRETRGLF | 9 | 9 | 100 |

## DISCUSSION

In this study, we demonstrated that our custom targeted genomic sequencing method can be used to effectively characterize IAV genomic material in wetlands sediment. All segments of the IAV genome were detected (Figure 1), and diverse HA and NA subtypes were observed (Figure 3). Multiple HA and NA subtypes were frequently detected in the same specimen (Figure 3), highlighting the advantages of environmental surveillance. The diversity of subtypes we observed showed that the custom probe panel designed for this study is broadly inclusive of diverse AIVs. It also revealed high HA and NA subtype richness among wild bird communities in the wetlands visited during the study period.

This method succeeded in recovering IAV genome fragments from specimens with low abundance of viral genomic material (Figure 2). Significant negative monotonic relationships were observed between screening RT-qPCR Ct values, the number of IAV genomic fragments recovered, and the number of IAV genome segments detected in a specimen (Figure 2). The practical implication of these results is that specimens with lower screening RT-qPCR Ct values (*i.e.* higher abundance of viral genomic material) should be prioritized when sequencing capacity is limited.

This method’s ability to recover IAV genome fragments from specimens with low abundance of viral genomic materials means that detections of particular segments or subtypes might rely on the recovery of limited numbers of genome fragments. This study demonstrated that these detections are credible. Even when the number of fragments detected in a specimen were limited, those fragments were described by hundreds to thousands of read pairs (Figure 4A). Furthermore, there was no evidence of cross-contamination in the 6 independent negative controls (Table S3), and chimeras and index hops were rare in data processed by HopDropper. The lack of evidence for false detections in this study reflects several method design choices that were made to increase confidence in results. IAV material is not used as positive control material so it cannot contaminate specimens. Negative controls contain sufficient background material to provide sensitive detection of low-level cross-contamination. UMIs are used during library construction to enable effective chimeric artefact removal.

This study also highlighted that the incomplete and fragmentary nature of IAV genomic material recovered from these specimens is a constraint of using wetlands sediment for genomics-based AIV surveillance. Only 20% of specimens had fragments from all 8 IAV genome segments recovered by probe capture-based targeted genomic sequencing (Figure 1C). Most fragments were short and only covered small regions of the IAV genome segment from which they originated (Figure 5).

Some longer fragments were recovered, but only up to 300 nucleotides were sequenced from each end of these fragments. This is an important trade-off for this method: Illumina short read platforms may leave the middles of longer fragments undescribed, but their high single-read accuracy and paired-end chemistry are instrumental for UMI-based single-fragment resolution and chimeric artefact removal. This trade-off seems prudent when enrichment and amplification are necessary to sequence fragmentary genomic material originating from multiple viruses in complex, challenging environmental specimens. If sequencing further along fragments is desired, paired-end sequencing runs could be performed with asymmetrical read lengths, *e.g.* 25 cycles for one read (to capture the UMI on that end of the molecule) followed by 575 cycles for the other read (to sequence further along the fragment)^33^. Alternatively, data generated by this method could be used to identify libraries containing long fragments of particularly high interest; these libraries could then be individually reflexed to a long-read platform.

In many applications, it is routine and appropriate to assemble fragmentary sequences into larger contigs. Full genomes might be instrumental for comparing genetic similarity between strains or constructing trees for phylodynamic analyses. The results from this study suggest that assembling the fragments recovered from sediment is not prudent. Genomic material from multiple viruses was often present in a single specimen (Figure 3C). Thus, assembling fragments may combine sequences originating from different viruses and create fictitious genomes. Each distinct fragment should be analyzed independently, and these fragments must be the unit of analysis for surveillance.

Fortunately, these distinct IAV genome fragments recovered from sediment provide useful information that can address practical surveillance questions. High quality alignments to well-characterized reference sequences were obtained (Figure S2). This allowed qualitative characterization of the potential host range and geographic origin of the IAVs in these specimens with the FindFlu tool (Figure 6, Figure 7, and Figure 8). This would allow the detection of viruses resembling those that have already spilled over into poultry and/or humans. It would also allow the detection of viral incursions from regions where pathogenic strains are known to circulate.

We also used FindFlu to identify the lineage and clade of recovered H5 fragments (Figure 8). This would allow the direct identification of HPAI gs/Gd lineage viruses, especially with the broad coverage of gs/Gd viruses provided by our custom probe panel (Table S2). Unfortunately, this study was unable to explicitly demonstrate detection of gs/Gd H5 viruses in sediment. No HPAI outbreaks occurred in the study region during the study period. A portion of specimen collection coincided with the arrival of a clade 2.3.4.4 gs/Gd lineage H5N1 virus in North America, but this strain did not arrive in the region where the study was conducted until April 2022^34,35^, three months after specimen collected had ended. Consequently, all H5 fragments detected originated from low pathogenicity, American H5 lineages that commonly circulate in waterfowl and shorebirds (Figure 6, Figure 7, Figure 8, and Figure 9).

While recovered H5 fragments could not be directly analyzed phylogenetically, we were able to infer their phylogenetic context by alignment to a proxy tree. Based on the positions in this tree of: 1) the prototypical gs/Gd sequence from 1996, 2) contemporary gs/Gd clades, and 3) the branch to which the recovered H5 fragments aligned, we surmised that the H5 viruses detected in these specimens were separated from gs/Gd viruses by several decades of evolution. A more precise estimate could be reached with phylodynamic modeling and a molecular clock, but this was beyond the scope of this study. Nonetheless, this demonstrates how fragmentary, incomplete sequences recovered from sediment could be incorporated into future phylodynamic analyses.

The results from this study highlight some interpretation challenges that might arise when using genomic data for surveillance. Notably, small minorities of fragments had their best matches to IAVs that have previously infected poultry and humans (Figure 4.8), but the level of risk implied by these detections was not clear. HPAI phenotypes are typically associated with specific HA subtypes, but none of these fragments originated from HA segments. Instead, they originated from NA and internal segments. These fragments may have belonged to low-risk strains that had acquired NA or internal segments from high-risk strains through re-assortment.

In these re-assortment scenarios, the segments acquired from spillover strains may not have been those that encoded the spillover phenotype. Another possibility is that these fragments merely originated from regions of IAV genome segments that are well-conserved between lineages circulating among different hosts. Another possibility to consider is that these viruses where not truly the closest matches for these genome fragments; better matching viruses from wild birds may have been absent from the bioinformatic database because viral diversity in wild birds is undersampled and incompletely characterized.

These interpretation challenges are not unique to probe capture methodology or environmental sampling, however. All genomic surveillance must contend with inferring viral phenotype and spillover risk from genotype, sequence similarity, and phylogenetic context. That is why this study assessed pathogenicity more directly by interrogating genetic markers of virulence on recovered fragments. We focused on H5 cleavage sites and corroborated the presence of low-pathogenicity H5 strains by detecting only canonical monobasic cleavage site motifs (Table 1). This same concept could be applied to other phenotypic markers of virulence and host range, however^36,37,38,39^.

The method presented in this study is flexible, and it could accommodate RNA extracted from various types of specimens. This expands its use to animal-based AIV surveillance as a culture-free method for direct sequencing of IAVs in bird swabs. This would avoid the extensive biocontainment infrastructure required for culturing suspected HPAIs, and it would be useful for sequencing swabs with low viral loads that fail conventional whole genome sequencing methods. Sequencing of wetlands sediment and bird swabs with this method would be easily scaled and parallelized; sediment and swab specimens could be processed simultaneously on the same library construction plates, captured in the same reaction, and sequenced on the same run, thereby increasing throughput and decreasing cost per specimen.

The One Health design of our custom probe panel further expands the types of specimens that could be assayed to include clinical specimens from other animals (*e.g.* swine and humans) as well as diverse environmental specimens (*e.g.* material from swine barns and agricultural fairs, filtered air from building HVAC systems, and municipal wastewater)^40,41,42,43,44^. Additionally, these specimens often contain other pathogens of importance to agriculture and public health, and probe panels could be expanded for simultaneous multi-pathogen detection^45,46,47,48^. In this way, the probe capture-based targeted genomic sequencing method demonstrated here could provide a powerful general-purpose tool for pathogen surveillance.

## MATERIALS AND METHODS

### Specimen collection

Sediment specimens were collected from 22 wetlands across the Metro Vancouver and Lower Mainland region of British Columbia, Canada. Superficial sediment was collected from twenty separated sites at each wetland, providing a total of 440 specimens for the study. Specimen collection occurred between October 6, 2021, and January 17, 2022. All 20 sites in a wetland were sampled on the same day. Wetland locations were selected in consultation with local biologists to determine areas that were likely to have high abundance of wild waterfowl. Within a wetland, sampling locations were selected to maximize potential of use by waterfowl (*e.g.* evidence on shoreline of recent waterfowl usage), ease of access to submerged sediment (*e.g.* water depth of less than 0.5 m), and to represent as much of the spatial extent of the wetland as possible. Sampling locations within a wetland were separated by 2 m or more. At each sampling location, biologists walked 1 to 2 m into the water and scooped the superficial layers of sediment into a 120 mL sterile urine collection container while wearing sterile nitrile gloves. Environmental data was then collected at each sampling location, including the geographic coordinates, an estimated water depth, water pH, water salinity, water temperature, and the presence or absence of fresh waterfowl feces at the shoreline.

Containers of sediment were brought back to the lab and kept at 4 °C until pre-processing could occur. During pre-processing, excess water was decanted, and large chunks of organic debris (*e.g.* leaves, plant roots, and rocks) were removed. The remaining material was manually stirred with a sterile metal scoopula to homogenize it, then 10 to 12 mL of the remaining material was placed into a 15 mL conical tube. The outsides of the tubes were wiped clean, disinfected with a 10% bleach solution, and then placed into a -80 °C freezer until RNA extraction.

### Total RNA extraction from sediment specimens and RT-qPCR screening for IAV genomic material

Total RNA was extracted from 435 of 440 total sediment specimens collected for this study (5 specimens had insufficient sediment for extraction). Total RNA was extracted from 2 g of sediment using the Qiagen RNeasy PowerSoil Total RNA kit (#12866). A chloroform extraction was added to the manufacturer’s protocol to remove additional PCR inhibitors. After the phenol:chloroform:isoamyl alcohol (pH 6.5-8.0) extraction step in the manufacturer’s protocol, the aqueous phase was transferred to a new container then mixed with an equal volume of chloroform. Phases were separated by centrifugation, then this chloroform extraction was repeated on the aqueous phase. The manufacturer’s protocol was resumed after the second chloroform extraction. RNA was eluted in 30 μL of nuclease-free water and stored at -80 °C.

IAV genomic material was detected by RT-qPCR targeting the matrix (M) segment (Table 2)^49^. 25 μL reactions were prepared with Applied Biosystems AgPath-ID One-Step RT PCR reagents (#4387391), 400 nM each of primers M52C and M253R (Table 2), 120 nM of the FAM-labeled probe M96C (Table 2), 3 μg of New England BioLabs T4 Gene 32 Protein (#M0300), and 2 μL of RNA extracted from sediment specimens. Reactions were incubated with the following cycling conditions: 1 cycle of 45 °C for 10 min; 1 cycle of 95 °C for 10 min; 45 cycles of 95 °C for 15 s followed by 60 °C for 60 s. Reactions were run on an Applied Biosystems 7500 Fast Real-Time PCR System using a fixed critical threshold of 0.05 for all reactions. Following common clinical practice, a Ct value of 40 was selected as the cut-off for specimen positivity. Screening RT-qPCR reactions were allowed to proceed for an additional 5 cycles, however, to identify suspect-positive specimens and assess their value for surveillance. Specimens were called suspect-positive if they had Ct values greater than 40 or if their amplification curves trended towards the critical threshold in the final PCR cycle.

**Table 2:**
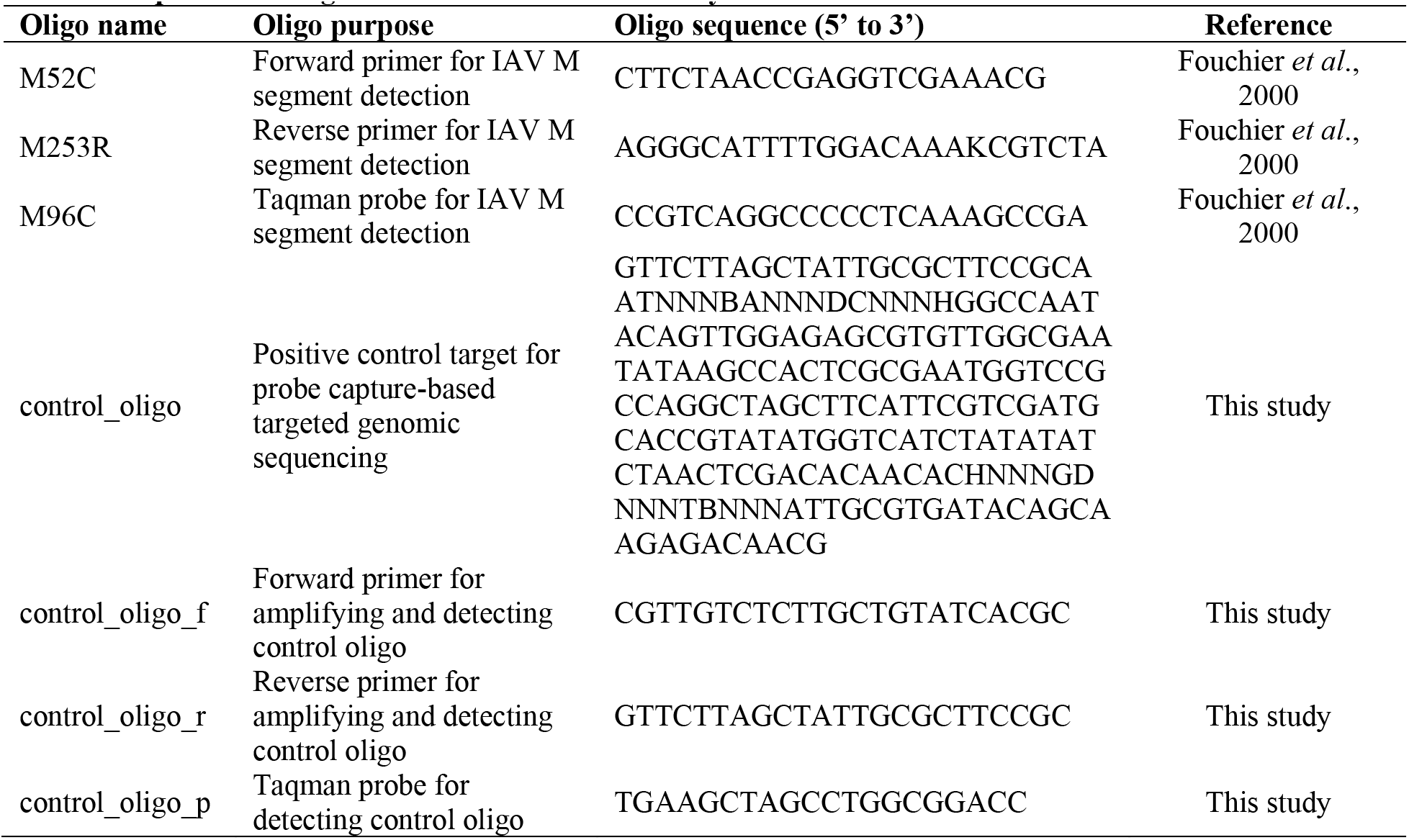
Sequences of oligonucleotides used in this study.

### cDNA synthesis and library construction

Double-stranded cDNA was prepared from 11 μL of undiluted RNA using the Invitrogen SuperScript IV First-Strand Synthesis System (#18091200) and the Invitrogen Second Strand cDNA Synthesis Kit (#A48571). First and second strand synthesis were both performed according to the manufacturer’s protocols, then purified using 1.8X Agencourt AMPure XP-PCR Purification Beads (#A63881). Sequencing libraries were prepared from the total volume of purified cDNA using the Integrated DNA Technologies xGen cfDNA & FFPE DNA Library Preparation Kit (#10006202) according to the manufacturer’s protocol. Libraries were barcoded using the xGen UDI Primers Plate 1 (#10005922) with 15 cycles of PCR. Following barcoding PCRs, libraries were purified with 1.3X Agencourt AMPure XP-PCR Purification Beads (#A63881) then eluted in 30 μL of nuclease-free water.

Libraries were prepared in batches of 15 sediment specimens and 1 batch control specimen. Sediment specimens were randomly assigned to 6 batches. All specimens in the same batch were prepared on the same reaction plates and from the same reagent master mixes. Batch controls were composed of 500 ng of Invitrogen Universal Human Reference RNA (#QS0639) spiked with 40,000 copies of double-stranded control oligo. The sequence of the control oligo was generated randomly (Table 2), then it was synthesized as an ssDNA Ultramer DNA Oligo by Integrated DNA Technologies (Coralville, Iowa, United States of America). Single-stranded control oligo was amplified by PCR as follows. 50 μL reactions were prepared with New England BioLabs NEBNext Ultra II Q5 Master Mix (#M0544), 1 μM of each control oligo amplification primer (Table 2), and 20 million copies of single-stranded control oligo Ultramer as template. Reactions were incubated with the following cycling conditions: 1 cycle of 98 °C for 30 s; 10 cycles of 98 °C for 15 s followed by 65 °C for 75 s; 1 cycle of 65 °C for 10 min. After amplification, double-stranded control oligo PCR products were purified using 1.2X Agencourt AMPure XP-PCR Purification Beads (#A63881) then eluted in 25 μL of nuclease-free water.

To spike batch controls with the specified copies of double-stranded control oligo, the molarity of the purified double-stranded control oligo PCR product was determined by qPCR. 20 μL reactions were prepared with New England BioLabs Luna Universal Probe qPCR Master Mix (#M3004), 250 nM of each control oligo amplification primer (Table 2), 250 nM of FAM-labeled control oligo detection probe (Table 2), and 2 μL of purified double-stranded control oligo PCR product. Reactions were run on an Applied Biosystems 7500 Fast Real-Time PCR System with the following cycling conditions: 1 cycle of 95 °C for 60 s; 40 cycles of 95 °C for 15 s followed by 60 °C for 45 s. A dilution series of the single-stranded control oligo Ultramer stock was used as a standard curve for quantification.

Enrichment of control oligos in batch control specimens functioned as a positive control for library construction and probe capture. Absence of control oligos in sediment specimens following index hop removal (described below) functioned as a negative control for reagent contamination and cross-contamination between specimens. Absence of IAV fragments in batch control specimens also functioned as a negative control in the same way.

### One Health IAV probe panel design

IAV genome segment sequences were downloaded from the Influenza Research Database (www.fludb.org)^50^ on December 9, 2021. Sequences were limited to those marked as complete from avian, swine, and human hosts. In total 531,526 IAV genome segment nucleotide sequences were obtained. Separate sub-panels were designed for each IAV genome segment as follows. First, all reference sequences representing a segment were clustered at 99% nucleotide identity using VSEARCH cluster_fast (v2.21.0) without masking (- qmask none)^51^. Cluster centroids were used as the input design space for ProbeTools *makeprobes* (v0.1.9) using batch sizes of 10 probes (-b 10), probe length of 120 nucleotides (-k 120), and a coverage endpoint of 95% (-c 95)^45^. Sub-panels for each IAV genome segment were combined to create the final panel. 10 additional probes with randomly generated sequences were added for capturing synthetic spike-in control oligos, although only one of these synthetic controls was used in this study (described above). The final panel contained 9,380 probes (sequences provided in Supplemental Material 1). ProbeTools *capture* and *stats* (v0.1.9) were used to confirm extensive coverage by the final panel of reference sequences in the design space (Table S1 and Table S2). The final panel was synthesized by Twist Biosciences (San Francisco, California, United States of America) with 0.02 fmol of each probe per reaction.

### Library pooling, hybridization probe capture, and genomic sequencing

dsDNA concentration was measure for each library using the Invitrogen Qubit dsDNA Broad Range kit (#Q32851) on the Invitrogen Qubit 4 Fluorometer. 300 ng of each library was pooled together, then two independent capture replicates were performed on aliquots of the pool. For each capture replicate, 2 aliquots of 4 μg of the pool were captured separately. After this first capture, they were combined and subjected to an additional capture for further enrichment of IAV genomic material. This means that 8 μg of library pool was enriched for each independent capture replicate and 16 μg of library pool was enriched in total for the whole study.

Pooled library material was completely evaporated in a vacuum oven at 50 °C and -20 mm Hg, then hybridization reactions were prepared with our custom One Health IAV panel (described above), Twist Universal Blockers (#100578), and Twist Hybridization Reagents (#104178) according to the manufacturer’s protocol. Hybridization reactions were incubated at 70 °C for 16 hours then washed with Twist Wash Buffers (#104178). Washing was performed according to the manufacturer’s protocol except the streptavidin bead slurry was resuspended in 22.5 μL of nuclease-free water instead of 45 μL prior to post-capture PCR. Post-capture PCR was performed on the total volume of bead slurry using Twist Equinox Library Amp Mix (#104178). Reactions were prepared and incubated according to the manufacturer’s protocol with 15 cycles of amplification. Following post-capture PCR, reactions were purified with the included DNA Purification Beads according to the manufacturer’s protocol. Purified captured library pools were eluted in 30 μL of nuclease-free water.

Molarity of double-captured library pools was determined using the New England BioLabs NEBNext Library Quant Kit for Illumina (#E7630). Double-captured library pools were also run on the Agilent TapeStation 2200 device using Agilent D1000 ScreenTape (#5067-5582) and D1000 reagents (#5067-5583) to obtain the peak fragment size, which was used to adjust molarity. 15 pmol of double-captured library pool was sequenced on an Illumina MiSeq instrument using MiSeq v3 600 cycle reagent kits (#MS-102-3003) to generate 2 x 300 cycle paired end reads. Each independent capture replicate was sequenced on its own run. The following adapter sequences were provided in the MiSeq sample sheet for on-instrument trimming: AGATCGGAAGAGCACACGTCTGAACTCCAGTCA and AGATCGGAAGAGCGTCGTGTAGGGAAAGAGTGT.

### Chimera and index hop removal and generation of consensus sequences for distinct fragments

Each MiSeq run was separately analyzed with HopDropper (v1.0.0) (https://github.com/KevinKuchinski/HopDropper). All FASTQ files generated in the run were analyzed, including sediment specimen libraries, control specimen libraries, and Undetermined libraries. 14-nucleotide intrinsic UMIs and 8-nucleotide extrinsic UMIs were assigned to each read, and extrinsic UMIs were limited to the 32 indices provided with the Integrated DNA Technologies xGen cfDNA & FFPE DNA Library Preparation Kit (#10006202). Fragments and their read pairs were only outputted if their UMI pair was observed at least 30 times. Fragment end consensus sequences were generated by sub-sampling up to 200 read pairs from each fragment. HopDropper defaults were used for other parameters.

### Identification and characterization of influenza A virus genome fragments

Fragment end consensus sequences generated by HopDropper were analyzed by FindFlu (v0.0.8) (https://github.com/KevinKuchinski/FindFlu). The FindFlu database used for this study was comprised of all complete segment nucleotide sequences in the Influenza Research Database (www.fludb.org) from avian, swine, and human hosts on October 11, 2022. IAV reference sequence were further filtered by length as follows: between 2,260 and 2,360 nucleotides for PB2 and PB1 segment sequences, between 2,120 and 2,250 nucleotides for PA segment sequences, between 1,650 and 1,800 nucleotides for HA segment sequences, between 1,480 and 1,580 nucleotides for NP segment sequences, between 1,250 and 1,560 nucleotides for NA segment sequences, between 975 and 1,030 nucleotides for M segment sequences, and between 815 and 900 nucleotides for NS segment sequences. The final database contained 169,098 avian-origin sequences, 70,918 swine-origin sequences, and 315,348 human-origin sequences (555,364 total sequences). IAV fragments from both probe capture replicates were combined for analyses in this study, except for analyses where probe capture replicates were explicitly considered separately. All fragment counts were based on UMI pair to ensure that IAV fragments were not double-counted if they were enriched in both probe capture replicates.

The FindFlu fragment report provided the following for each IAV fragment: segment, subtype, number of copies sequenced, fragment length, segment coverage, alignment identity, and alignment query coverage. The FindFlu host, country, and H5 clade reports were used to calculate the percentage of IAV fragments having their best matches to reference sequences from various host species, collection countries, and H5 clades. In cases where IAV fragments had multiple best-matching reference sequences with multiple host/country/H5 clade annotations, each different host/country/H5 clade value observed was proportionally allocated a fraction of a fragment (1/*n* where *n* was the number of best-matching reference sequences the fragment had).

### Phylogenetic analysis of H5 fragments

Recent H5 segment reference sequences were downloaded from the Influenza Research Database (www.fludb.org)^50^. All available complete H5 segment nucleotide sequences collected from 2018 onwards were downloaded on November 6, 2022. The prototypical goose/Guangdong/96 lineage HA sequence (GenBank accession NC_007362) was also included to represent Clade 0. A multiple sequence alignment was performed on the resulting collection of 147 H5 reference sequences using CLUSTAL W (v2.1)^52^. A maximum likelihood phylogenetic tree was constructed from the multiple sequence alignment and bootstrapped 100 times using PHYML (v3.3.20211231)^53^. The resulting tree was analyzed and visualized with the ETE3 package (v3.1.2) in Python (v3.9.12)^54^. Outlying branches were trimmed if their length exceeded 3 standard deviations of the mean branch length. For visual clarity, monophyletic groups of similar leaves were collapsed into a single leaf if all leaves were less than 0.025 substitutions/site from their common ancestral node. The length of the replacement leaf’s branch was set to the mean branch length of the collapsed leaves.

The best-matching reference sequences for each H5 fragment were determined as follows. The H5 fragment end consensus sequences generated by HopDropper were aligned to the H5 reference sequences composing the tree using blastn (v2.13.0)^55^. A combined bitscore was generated for each fragment-reference sequence combination by adding together the bitscores from both fragment end consensus sequence alignments against that reference sequence. Each fragment’s best-matching reference sequences were those with which it had its maximum combined bitscore.

The percentage of H5 fragments having their best match to each reference sequence composing the tree was calculated as follows. The number of H5 fragments having their best match to a reference sequence was divided by the total number of H5 fragments then multiplied by 100. In cases where an H5 fragment had multiple best matches, that fragment was counted as 1/*n* of a fragment for each of their best matches, where *n* was that fragment’s number of best matches. When similar tree leaves were collapsed into a single leaf for visual clarity, the replacement leaf’s percentage of H5 fragments having their best match to it was determined by summing the percentages of its constituent leaves.

### H5 cleavage site characterization

H5 fragment end consensus sequences generated by HopDropper were translated and aligned to the prototypical goose/Guangdong/96 lineage HA amino acid sequence (GenBank accession NC_007362) using blastx (v2.13.0)^55^. Only the best alignments (by bitscore) were retained for each fragment end consensus sequence. The position of each fragment end consensus sequence in the goose/Guangdong H5 amino acid sequence was determined from the alignment subject start and subject end coordinates. Fragment end consensus sequences containing the HA cleavage site were identified by finding fragments that spanned the coordinates 336 and 348. HA cleavage site motifs were then extracted from the aligned, translated sequences.

## ACKNOWLEDGEMENTS

We would like to thank all the laboratories worldwide who have submitted genomic sequences to the Influenza Research Database. We would also like to thank the Public Health Laboratory at the British Columbia Centre for Disease Control for maintaining laboratory space, RT-qPCR instruments, and the Illumina MiSeq sequencing platform used in this study. Ciara O’Higgins and Kristen Moffit at the BC Ministry of Agriculture and Food’s Animal Health Centre provided invaluable assistance with specimen processing. The EBE Environmental Consulting Inc. team’s dedication to collecting specimens in exceptionally challenging weather conditions was greatly appreciated.

## AUTHOR CONTRIBUTIONS

KK conceived the study, designed the custom panel of hybridization probes, developed laboratory methods and bioinformatic tools used to generate genomic data, analyzed and interpreted genomic data, and wrote the manuscript. MC conceived the study, developed the wetlands sampling strategy and sediment specimen collection protocol, oversaw specimen collection, and reviewed the manuscript. SM oversaw and troubleshot RNA extraction and RT-qPCR screening of sediment specimens and reviewed the manuscript. GC and MK assisted with troubleshooting and performed RNA extraction, RT-qPCR screening, library construction, and hybridization probe capture, and they reviewed the manuscript. CH conceived the study, secured funding, provided graduate-level supervision of MC, and reviewed the manuscript. NP conceived the study, secured funding, provided graduate-level supervision of KK, and reviewed the manuscript.

## DATA AVAILABILITY

Source code for HopDropper and FindFlu are available at https://github.com/KevinKuchinski/. Raw sequencing reads generated during this study are available from the NCBI Short Read Archive as part of BioProject PRJNA926989. Influenza A virus genome fragments recovered in this study (following HopDropper and FindFlu analysis, as described above) have been included as a supplemental FASTA file (Supplemental Material 2).

---

**Figure S1:**
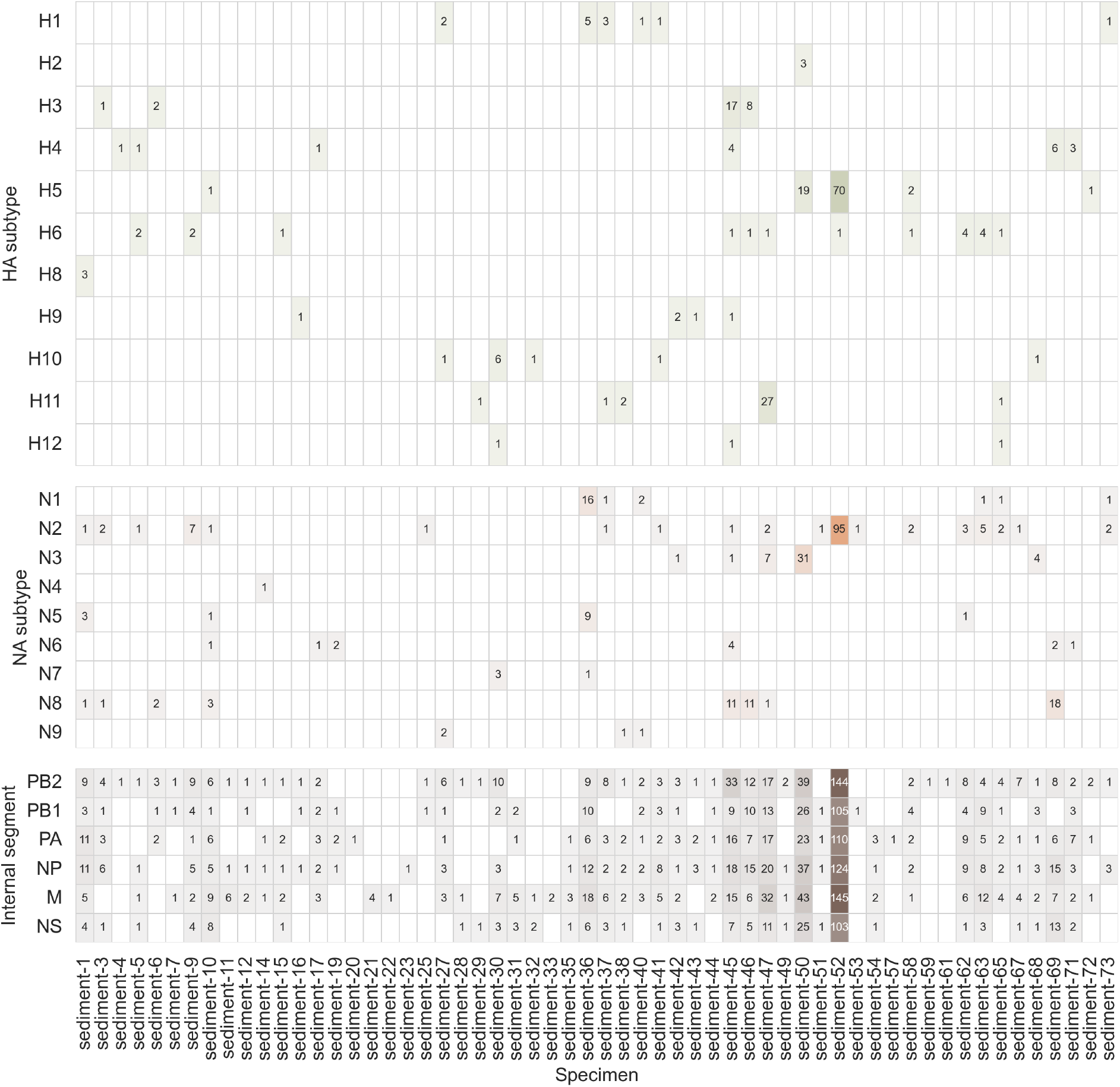
Number of influenza A virus genome fragments detected in wetlands sediment by probe capture-based targeted genomic sequencing. 2,312 fragments of influenza A virus (IAV) genome were recovered from 74 sediment specimens that tested positive for IAV genomic material by RT-qPCR. Numbers inside cells indicate the number of IAV fragments originating from a particular segment/subtype that were recovered from the corresponding specimen. Only specimens that contained IAV fragments were plotted.

**Figure S2:**
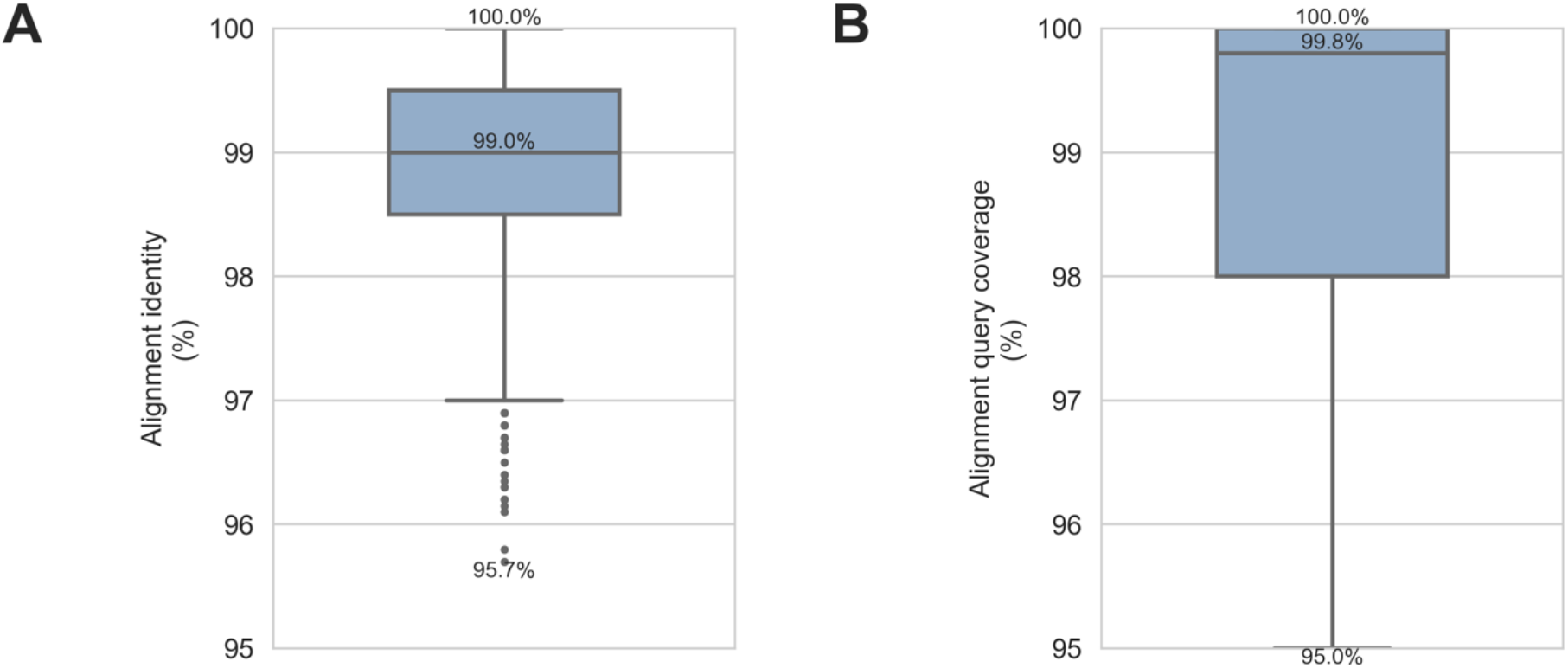
Influenza A virus genome fragments recovered from wetlands sediment were highly similar to reference sequences used by FindFlu to characterize viruses. FindFlu aligned recovered influenza A virus (IAV) genome fragments to a database of 555,364 IAV reference sequences (collected globally from avian, swine, and human hosts). **A)** Nucleotide sequence identities were calculated by dividing the number of identical bases by the alignment length. **B)** Query sequence coverage values were calculated by dividing alignment lengths by query sequence lengths. The minimum, median, and maximum values in both distributions are indicated on the plots.

**Table S1:** Custom probe panel provides broadly inclusive coverage of influenza A virus reference sequences. The ProbeTools capture and stats modules were used to predict *in silico* how well this study’s custom panel of 9,380 probes covered 531,526 influenza A virus (IAV) reference sequences (collected globally from avian, swine, and human hosts). For each reference sequence, probe coverage was calculated as the number of nucleotide positions covered by at least one probe in the panel. The minimum, 5^th^ percentile, median, and maximum probe coverage values were reported for each segment, subtype, and host category.

| Segment | Subtype | Host | Reference sequences (#) | Minimum coverage (%) | Fifth percentile of coverage (%) | Median coverage (%) | Maximum coverage (%) |
| --- | --- | --- | --- | --- | --- | --- | --- |
| PB2 | n/a | avian | 19109 | 71.4 | 94.5 | 99.3 | 100.0 |
| PB2 | n/a | human | 34153 | 86.4 | 91.8 | 97.8 | 100.0 |
| PB2 | n/a | swine | 6767 | 73.0 | 95.6 | 99.1 | 100.0 |
| PB1 | n/a | avian | 18173 | 70.9 | 94.2 | 99.7 | 100.0 |
| PB1 | n/a | human | 28158 | 82.9 | 95.8 | 99.8 | 100.0 |
| PB1 | n/a | swine | 5939 | 78.7 | 94.6 | 99.7 | 100.0 |
| PA | n/a | avian | 19990 | 62.6 | 93.9 | 99.9 | 100.0 |
| PA | n/a | human | 34391 | 80.7 | 93.9 | 98.1 | 100.0 |
| PA | n/a | swine | 6919 | 78.0 | 94.9 | 99.7 | 100.0 |
| HA | all | avian | 26173 | 62.1 | 93.9 | 99.9 | 100.0 |
| HA | all | human | 49011 | 82.3 | 98.5 | 99.8 | 100.0 |
| HA | all | swine | 11342 | 67.9 | 95 | 99.9 | 100.0 |
| HA | H1 | avian | 916 | 73.3 | 91.6 | 98.1 | 100.0 |
| HA | H1 | human | 21844 | 82.3 | 98.5 | 100.0 | 100.0 |
| HA | H1 | swine | 7899 | 72.2 | 95.1 | 99.8 | 100.0 |
| HA | H2 | avian | 530 | 81.7 | 90 | 99.4 | 100.0 |
| HA | H2 | human | 95 | 93.6 | 98.8 | 99.6 | 100.0 |
| HA | H2 | swine | 2 | 97.5 | 97.6 | 98.1 | 98.7 |
| HA | H3 | avian | 2269 | 66.6 | 92.4 | 99.1 | 100.0 |
| HA | H3 | human | 26651 | 89.6 | 98.5 | 99.0 | 100.0 |
| HA | H3 | swine | 3374 | 67.9 | 94.7 | 99.9 | 100.0 |
| HA | H4 | avian | 1928 | 72.8 | 93.9 | 99.4 | 100.0 |
| HA | H4 | human | 1 | 100.0 | 100 | 100.0 | 100.0 |
| HA | H4 | swine | 5 | 97.3 | 97.3 | 99.4 | 100.0 |
| HA | H5 | avian | 5555 | 75.4 | 96.7 | 100.0 | 100.0 |
| HA | H5 | human | 246 | 94.2 | 98.5 | 100.0 | 100.0 |
| HA | H5 | swine | 33 | 98.8 | 99.4 | 100.0 | 100.0 |
| HA | H6 | avian | 1902 | 77.4 | 94.2 | 99.8 | 100.0 |
| HA | H6 | swine | 2 | 96.6 | 96.8 | 98.3 | 100.0 |
| HA | H7 | avian | 2367 | 70.6 | 93.8 | 100.0 | 100.0 |
| HA | H7 | human | 152 | 93.2 | 96.1 | 100.0 | 100.0 |
| HA | H7 | swine | 3 | 85.7 | 86.9 | 98.0 | 98.0 |
| HA | H8 | avian | 160 | 79.6 | 92.8 | 98.7 | 100.0 |
| HA | H9 | avian | 7364 | 73.1 | 96.2 | 100.0 | 100.0 |
| HA | H9 | human | 18 | 87.9 | 93.3 | 100.0 | 100.0 |
| HA | H9 | swine | 22 | 87.5 | 92.1 | 97.6 | 100.0 |
| HA | H10 | avian | 1231 | 81.8 | 92.8 | 99.9 | 100.0 |
| HA | H10 | human | 4 | 98.5 | 98.5 | 98.5 | 98.5 |
| HA | H10 | swine | 1 | 99.4 | 99.4 | 99.4 | 99.4 |
| HA | H11 | avian | 736 | 77.5 | 93.3 | 99.9 | 100.0 |
| HA | H11 | swine | 1 | 82.7 | 82.7 | 82.7 | 82.7 |
| HA | H12 | avian | 344 | 78.5 | 92.6 | 98.1 | 100.0 |
| HA | H13 | avian | 546 | 75.4 | 91 | 97.0 | 100.0 |
| HA | H14 | avian | 39 | 79.3 | 85 | 100.0 | 100.0 |
| HA | H15 | avian | 20 | 62.1 | 62.1 | 79.4 | 100.0 |
| HA | H16 | avian | 266 | 69.2 | 81.0 | 98.4 | 100.0 |
| NP | n/a | avian | 19897 | 67.9 | 94.7 | 99.5 | 100.0 |
| NP | n/a | human | 36140 | 71.8 | 93.7 | 96.7 | 100.0 |
| NP | n/a | swine | 7168 | 68.8 | 94.2 | 98.3 | 100.0 |
| NA | all | avian | 20401 | 63.5 | 93.2 | 99.7 | 100.0 |
| NA | all | human | 43304 | 74.7 | 98.6 | 100.0 | 100.0 |
| NA | all | swine | 12126 | 69.4 | 95.8 | 100.0 | 100.0 |
| NA | N1 | avian | 4446 | 78.8 | 95.0 | 98.8 | 100.0 |
| NA | N1 | human | 19974 | 74.7 | 98.5 | 99.9 | 100.0 |
| NA | N1 | swine | 4936 | 76.7 | 96.0 | 99.6 | 100.0 |
| NA | N2 | avian | 5341 | 79.9 | 94.7 | 99.8 | 100.0 |
| NA | N2 | human | 23200 | 86.9 | 99.7 | 100.0 | 100.0 |
| NA | N2 | swine | 7171 | 69.4 | 95.8 | 100.0 | 100.0 |
| NA | N3 | avian | 1681 | 74.8 | 91.7 | 98.5 | 100.0 |
| NA | N3 | human | 2 | 95.7 | 95.8 | 96.4 | 97.0 |
| NA | N3 | swine | 4 | 98.1 | 98.1 | 99.0 | 99.9 |
| NA | N4 | avian | 378 | 81.1 | 90.0 | 99.8 | 100.0 |
| NA | N4 | human | 1 | 96.3 | 96.3 | 96.3 | 96.3 |
| NA | N5 | avian | 567 | 77.7 | 89.7 | 99.2 | 100.0 |
| NA | N5 | swine | 1 | 94.3 | 94.3 | 94.3 | 94.3 |
| NA | N6 | avian | 2930 | 78.3 | 92.9 | 99.9 | 100.0 |
| NA | N6 | human | 4 | 100.0 | 100.0 | 100.0 | 100.0 |
| NA | N6 | swine | 8 | 93.6 | 93.7 | 99.9 | 100.0 |
| NA | N7 | avian | 1078 | 63.5 | 94.0 | 100.0 | 100.0 |
| NA | N7 | human | 4 | 97.5 | 97.5 | 98.2 | 98.8 |
| NA | N7 | swine | 1 | 74.1 | 74.1 | 74.1 | 74.1 |
| NA | N8 | avian | 2528 | 74.9 | 92.7 | 99.8 | 100.0 |
| NA | N8 | human | 5 | 99.8 | 99.8 | 100.0 | 100.0 |
| NA | N8 | swine | 3 | 89.2 | 89.2 | 89.2 | 97.0 |
| NA | N9 | avian | 1452 | 72.9 | 93.3 | 100.0 | 100.0 |
| NA | N9 | human | 114 | 95.5 | 98.1 | 100.0 | 100.0 |
| NA | N9 | swine | 2 | 100.0 | 100.0 | 100.0 | 100.0 |
| M | n/a | avian | 20263 | 77.0 | 92.7 | 100.0 | 100.0 |
| M | n/a | human | 40456 | 78.2 | 89.5 | 99.8 | 100.0 |
| M | n/a | swine | 9860 | 73.7 | 98.2 | 100.0 | 100.0 |
| NS | n/a | avian | 20810 | 66.5 | 95.4 | 99.8 | 100.0 |
| NS | n/a | human | 34027 | 76.1 | 98.3 | 100.0 | 100.0 |
| NS | n/a | swine | 6949 | 64.4 | 96.2 | 100.0 | 100.0 |

**Table S2:**
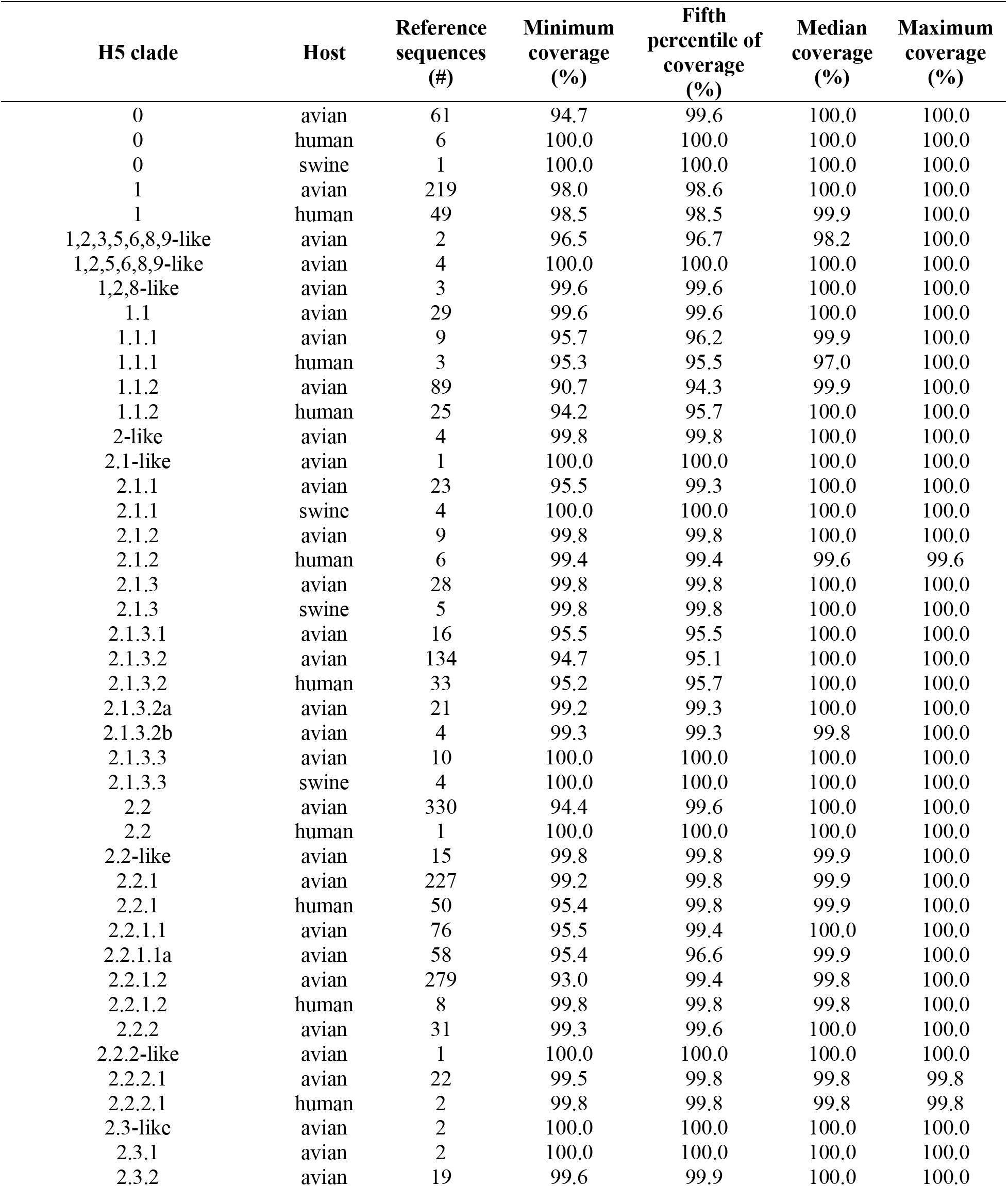
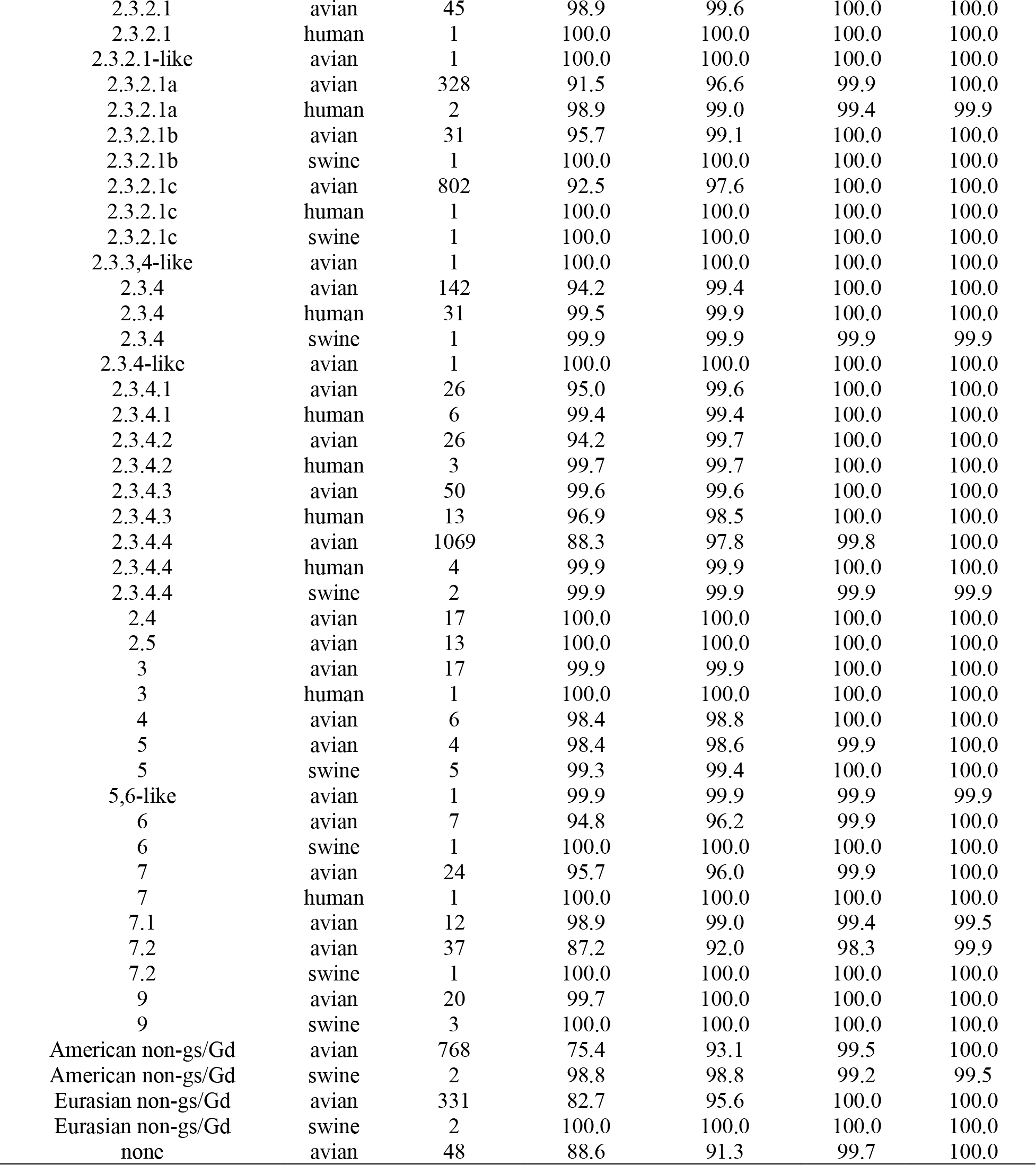
Custom probe panel provides broadly inclusive coverage of H5 subtype haemagglutinin segment reference sequences from diverse clades of the goose/Guangdong/96 lineage. The ProbeTools captures and stats modules were used to predict *in silico* how well this study’s custom panel of 9,380 probes covered 5,834 H5 subtype haemagglutinin (HA) segment reference sequences (collected globally from avian, swine, and human hosts). For each reference sequence, probe coverage was calculated as the number of nucleotide positions covered by at least one probe in the panel. The minimum, 5^th^ percentile, median, and maximum probe coverage values were reported for each segment, subtype, and host category.

**Table S3:** Probe capture and sequencing metrics. Sequencing libraries were prepared from 90 specimens that were positive (n=74) or suspect-positive (n=16) for influenza A virus (IAV) genomic material by RT-qPCR. After pooling libraries together, two independent probe captures were performed and sequenced separately (capture_1 and capture_2). FASTQ data was analyzed with HopDropper to identify distinct fragments of genomic material and remove chimeric artefacts (invalid reads). The number of total read pairs and valid read pairs was reported for each library by HopDropper. Fragment consensus sequences generated by HopDropper were analyzed by FindFlu to identify fragments of IAV genome. The number of IAV read pairs in each library was determined by summing the number of copies for each IAV fragment identified by FindFlu. For each library, the number of IAV read pairs was divided by the total number of read pairs to calculate total on-target rates. Valid on-target rates were calculated by dividing the number of IAV read pairs in a library by the number of valid read pairs in that library.

| Capture replicate | Specimen ID | Screening RT-qPCR result | Screening RT-qPCR Ct value | Total read pairs (#) | Valid read pairs (#) | IAV read pairs (#) | Total on-target (%) | Valid on-target (%) |
| --- | --- | --- | --- | --- | --- | --- | --- | --- |
| capture-1 | sediment-1 | Positive | 29.74 | 172839 | 123424 | 82372 | 47.7 | 66.7 |
| capture-2 | sediment-1 | Positive | 29.74 | 184143 | 125720 | 80859 | 43.9 | 64.3 |
| capture-1 | sediment-2 | Positive | 37.80 | 9816 | 0 | 0 | 0.0 | 0.0 |
| capture-2 | sediment-2 | Positive | 37.80 | 13078 | 0 | 0 | 0.0 | 0.0 |
| capture-1 | sediment-3 | Positive | 34.64 | 362356 | 239693 | 205456 | 56.7 | 85.7 |
| capture-2 | sediment-3 | Positive | 34.64 | 317087 | 212259 | 173528 | 54.7 | 81.8 |
| capture-1 | sediment-4 | Positive | 37.14 | 617838 | 415482 | 415482 | 67.2 | 100.0 |
| capture-2 | sediment-4 | Positive | 37.14 | 657293 | 454675 | 454675 | 69.2 | 100.0 |
| capture-1 | sediment-5 | Positive | 34.39 | 28854 | 16351 | 13818 | 47.9 | 84.5 |
| capture-2 | sediment-5 | Positive | 34.39 | 35749 | 21026 | 11936 | 33.4 | 56.8 |
| capture-1 | sediment-6 | Positive | 35.81 | 61294 | 35924 | 33763 | 55.1 | 94.0 |
| capture-2 | sediment-6 | Positive | 35.81 | 73277 | 42115 | 33868 | 46.2 | 80.4 |
| capture-1 | sediment-7 | Positive | 37.17 | 13249 | 5330 | 2047 | 15.5 | 38.4 |
| capture-2 | sediment-7 | Positive | 37.17 | 15571 | 3476 | 2221 | 14.3 | 63.9 |
| capture-1 | sediment-8 | Positive | 37.66 | 7102 | 0 | 0 | 0.0 | 0.0 |
| capture-2 | sediment-8 | Positive | 37.66 | 9909 | 0 | 0 | 0.0 | 0.0 |
| capture-1 | sediment-9 | Positive | 31.39 | 220956 | 150388 | 110129 | 49.8 | 73.2 |
| capture-2 | sediment-9 | Positive | 31.39 | 187552 | 129177 | 95429 | 50.9 | 73.9 |
| capture-1 | sediment-10 | Positive | 33.79 | 59842 | 37111 | 15779 | 26.4 | 42.5 |
| capture-2 | sediment-10 | Positive | 33.79 | 76291 | 46956 | 20288 | 26.6 | 43.2 |
| capture-1 | sediment-11 | Positive | 34.94 | 40338 | 21756 | 7707 | 19.1 | 35.4 |
| capture-2 | sediment-11 | Positive | 34.94 | 71886 | 43453 | 31076 | 43.2 | 71.5 |
| capture-1 | sediment-12 | Positive | 37.25 | 30905 | 16288 | 10776 | 34.9 | 66.2 |
| capture-2 | sediment-12 | Positive | 37.25 | 40646 | 21804 | 13470 | 33.1 | 61.8 |
| capture-1 | sediment-13 | Positive | 37.53 | 530 | 0 | 0 | 0.0 | 0.0 |
| capture-2 | sediment-13 | Positive | 37.53 | 642 | 0 | 0 | 0.0 | 0.0 |
| capture-1 | sediment-14 | Positive | 37.74 | 6038 | 1321 | 781 | 12.9 | 59.1 |
| capture-2 | sediment-14 | Positive | 37.74 | 11660 | 4015 | 2628 | 22.5 | 65.5 |
| capture-1 | sediment-15 | Positive | 36.97 | 33403 | 14578 | 5164 | 15.5 | 35.4 |
| capture-2 | sediment-15 | Positive | 36.97 | 40416 | 13794 | 3763 | 9.3 | 27.3 |
| capture-1 | sediment-16 | Positive | 39.90 | 18514 | 3857 | 1853 | 10.0 | 48.0 |
| capture-2 | sediment-16 | Positive | 39.90 | 28731 | 9055 | 0 | 0.0 | 0.0 |
| capture-1 | sediment-17 | Positive | 36.04 | 99126 | 61623 | 33540 | 33.8 | 54.4 |
| capture-2 | sediment-17 | Positive | 36.04 | 124105 | 81015 | 42909 | 34.6 | 53.0 |
| capture-1 | sediment-18 | Positive | 36.91 | 16335 | 759 | 0 | 0.0 | 0.0 |
| capture-2 | sediment-18 | Positive | 36.91 | 21078 | 0 | 0 | 0.0 | 0.0 |
| capture-1 | sediment-19 | Positive | 38.27 | 24676 | 8550 | 8397 | 34.0 | 98.2 |
| capture-2 | sediment-19 | Positive | 38.27 | 24111 | 6567 | 6567 | 27.2 | 100.0 |
| capture-1 | sediment-20 | Positive | 39.54 | 7927 | 652 | 0 | 0.0 | 0.0 |
| capture-2 | sediment-20 | Positive | 39.54 | 11547 | 1809 | 1809 | 15.7 | 100.0 |
| capture-1 | sediment-21 | Positive | 38.93 | 16819 | 3886 | 1191 | 7.1 | 30.6 |
| capture-2 | sediment-21 | Positive | 38.93 | 21911 | 4759 | 4084 | 18.6 | 85.8 |
| capture-1 | sediment-22 | Positive | 38.16 | 10044 | 0 | 0 | 0.0 | 0.0 |
| capture-2 | sediment-22 | Positive | 38.16 | 13722 | 148 | 148 | 1.1 | 100.0 |
| capture-1 | sediment-23 | Positive | 37.49 | 28942 | 15618 | 9076 | 31.4 | 58.1 |
| capture-2 | sediment-23 | Positive | 37.49 | 34369 | 18373 | 7740 | 22.5 | 42.1 |
| capture-1 | sediment-24 | Positive | 39.23 | 192 | 0 | 0 | 0.0 | 0.0 |
| capture-2 | sediment-24 | Positive | 39.23 | 205 | 0 | 0 | 0.0 | 0.0 |
| capture-1 | sediment-25 | Positive | 38.26 | 18810 | 9709 | 2416 | 12.8 | 24.9 |
| capture-2 | sediment-25 | Positive | 38.26 | 11653 | 3375 | 33 | 0.3 | 1.0 |
| capture-1 | sediment-26 | Positive | 37.92 | 17050 | 4401 | 0 | 0.0 | 0.0 |
| capture-2 | sediment-26 | Positive | 37.92 | 20050 | 3748 | 0 | 0.0 | 0.0 |
| capture-1 | sediment-27 | Positive | 35.76 | 48138 | 25237 | 9404 | 19.5 | 37.3 |
| capture-2 | sediment-27 | Positive | 35.76 | 51918 | 25737 | 6616 | 12.7 | 25.7 |
| capture-1 | sediment-28 | Positive | 37.75 | 42399 | 20550 | 13367 | 31.5 | 65.0 |
| capture-2 | sediment-28 | Positive | 37.75 | 45617 | 19371 | 9055 | 19.9 | 46.7 |
| capture-1 | sediment-29 | Positive | 38.79 | 19025 | 4609 | 4047 | 21.3 | 87.8 |
| capture-2 | sediment-29 | Positive | 38.79 | 23342 | 3603 | 529 | 2.3 | 14.7 |
| capture-1 | sediment-30 | Positive | 33.48 | 296600 | 200680 | 92475 | 31.2 | 46.1 |
| capture-2 | sediment-30 | Positive | 33.48 | 303294 | 208434 | 100689 | 33.2 | 48.3 |
| capture-1 | sediment-31 | Positive | 36.04 | 51796 | 30680 | 15993 | 30.9 | 52.1 |
| capture-2 | sediment-31 | Positive | 36.04 | 53945 | 31069 | 24441 | 45.3 | 78.7 |
| capture-1 | sediment-32 | Positive | 34.38 | 118096 | 76146 | 60574 | 51.3 | 79.5 |
| capture-2 | sediment-32 | Positive | 34.38 | 107319 | 69349 | 49813 | 46.4 | 71.8 |
| capture-1 | sediment-33 | Positive | 37.00 | 12889 | 2179 | 0 | 0.0 | 0.0 |
| capture-2 | sediment-33 | Positive | 37.00 | 18721 | 4587 | 4203 | 22.5 | 91.6 |
| capture-1 | sediment-34 | Positive | 38.66 | 7925 | 0 | 0 | 0.0 | 0.0 |
| capture-2 | sediment-34 | Positive | 38.66 | 10580 | 146 | 0 | 0.0 | 0.0 |
| capture-1 | sediment-35 | Positive | 35.24 | 96018 | 57214 | 35142 | 36.6 | 61.4 |
| capture-2 | sediment-35 | Positive | 35.24 | 99988 | 49410 | 29437 | 29.4 | 59.6 |
| capture-1 | sediment-36 | Positive | 34.21 | 318299 | 207819 | 106905 | 33.6 | 51.4 |
| capture-2 | sediment-36 | Positive | 34.21 | 297011 | 189799 | 81687 | 27.5 | 43.0 |
| capture-1 | sediment-37 | Positive | 35.14 | 59164 | 30848 | 11874 | 20.1 | 38.5 |
| capture-2 | sediment-37 | Positive | 35.14 | 59973 | 31689 | 17038 | 28.4 | 53.8 |
| capture-1 | sediment-38 | Positive | 39.48 | 57390 | 23454 | 17437 | 30.4 | 74.3 |
| capture-2 | sediment-38 | Positive | 39.48 | 56215 | 20925 | 7968 | 14.2 | 38.1 |
| capture-1 | sediment-39 | Positive | 37.53 | 14108 | 0 | 0 | 0.0 | 0.0 |
| capture-2 | sediment-39 | Positive | 37.53 | 19305 | 0 | 0 | 0.0 | 0.0 |
| capture-1 | sediment-40 | Positive | 35.43 | 52572 | 29150 | 13058 | 24.8 | 44.8 |
| capture-2 | sediment-40 | Positive | 35.43 | 53874 | 22204 | 7616 | 14.1 | 34.3 |
| capture-1 | sediment-41 | Positive | 36.85 | 49970 | 29206 | 17826 | 35.7 | 61.0 |
| capture-2 | sediment-41 | Positive | 36.85 | 49103 | 26999 | 9784 | 19.9 | 36.2 |
| capture-1 | sediment-42 | Positive | 34.88 | 123004 | 80528 | 60880 | 49.5 | 75.6 |
| capture-2 | sediment-42 | Positive | 34.88 | 125657 | 81782 | 48570 | 38.7 | 59.4 |
| capture-1 | sediment-43 | Positive | 36.67 | 26365 | 8973 | 7365 | 27.9 | 82.1 |
| capture-2 | sediment-43 | Positive | 36.67 | 24860 | 6651 | 3996 | 16.1 | 60.1 |
| capture-1 | sediment-44 | Positive | 37.97 | 64481 | 39264 | 20760 | 32.2 | 52.9 |
| capture-2 | sediment-44 | Positive | 37.97 | 62189 | 35994 | 31542 | 50.7 | 87.6 |
| capture-1 | sediment-45 | Positive | 34.01 | 327306 | 212395 | 87909 | 26.9 | 41.4 |
| capture-2 | sediment-45 | Positive | 34.01 | 322004 | 215898 | 83987 | 26.1 | 38.9 |
| capture-1 | sediment-46 | Positive | 33.75 | 281204 | 178445 | 90685 | 32.2 | 50.8 |
| capture-2 | sediment-46 | Positive | 33.75 | 318599 | 209473 | 125384 | 39.4 | 59.9 |
| capture-1 | sediment-47 | Positive | 31.00 | 293899 | 178571 | 71589 | 24.4 | 40.1 |
| capture-2 | sediment-47 | Positive | 31.00 | 335264 | 205977 | 54692 | 16.3 | 26.6 |
| capture-1 | sediment-48 | Positive | 37.37 | 12068 | 1220 | 0 | 0.0 | 0.0 |
| capture-2 | sediment-48 | Positive | 37.37 | 15492 | 1243 | 0 | 0.0 | 0.0 |
| capture-1 | sediment-49 | Positive | 35.74 | 20204 | 6623 | 750 | 3.7 | 11.3 |
| capture-2 | sediment-49 | Positive | 35.74 | 25349 | 6566 | 3786 | 14.9 | 57.7 |
| capture-1 | sediment-50 | Positive | 30.51 | 434665 | 265394 | 122267 | 28.1 | 46.1 |
| capture-2 | sediment-50 | Positive | 30.51 | 399178 | 263511 | 111366 | 27.9 | 42.3 |
| capture-1 | sediment-51 | Positive | 38.78 | 32628 | 14046 | 10786 | 33.1 | 76.8 |
| capture-2 | sediment-51 | Positive | 38.78 | 45528 | 21216 | 17323 | 38.0 | 81.7 |
| capture-1 | sediment-52 | Positive | 27.86 | 2845588 | 1869576 | 1163089 | 40.9 | 62.2 |
| capture-2 | sediment-52 | Positive | 27.86 | 2879923 | 1924536 | 1195950 | 41.5 | 62.1 |
| capture-1 | sediment-53 | Positive | 37.85 | 12915 | 3572 | 2756 | 21.3 | 77.2 |
| capture-2 | sediment-53 | Positive | 37.85 | 17538 | 3878 | 1405 | 8.0 | 36.2 |
| capture-1 | sediment-54 | Positive | 38.26 | 70353 | 47014 | 37520 | 53.3 | 79.8 |
| capture-2 | sediment-54 | Positive | 38.26 | 101699 | 67246 | 53801 | 52.9 | 80.0 |
| capture-1 | sediment-55 | Positive | 37.32 | 24 | 0 | 0 | 0.0 | 0.0 |
| capture-2 | sediment-55 | Positive | 37.32 | 20 | 0 | 0 | 0.0 | 0.0 |
| capture-1 | sediment-56 | Positive | 39.89 | 8834 | 0 | 0 | 0.0 | 0.0 |
| capture-2 | sediment-56 | Positive | 39.89 | 11661 | 0 | 0 | 0.0 | 0.0 |
| capture-1 | sediment-57 | Positive | 36.32 | 21516 | 0 | 0 | 0.0 | 0.0 |
| capture-2 | sediment-57 | Positive | 36.32 | 31734 | 7919 | 7919 | 25.0 | 100.0 |
| capture-1 | sediment-58 | Positive | 36.83 | 35728 | 15287 | 3075 | 8.6 | 20.1 |
| capture-2 | sediment-58 | Positive | 36.83 | 32223 | 12298 | 4398 | 13.6 | 35.8 |
| capture-1 | sediment-59 | Positive | 33.37 | 9519 | 999 | 493 | 5.2 | 49.3 |
| capture-2 | sediment-59 | Positive | 33.37 | 11220 | 161 | 0 | 0.0 | 0.0 |
| capture-1 | sediment-60 | Positive | 37.16 | 3027 | 0 | 0 | 0.0 | 0.0 |
| capture-2 | sediment-60 | Positive | 37.16 | 4094 | 0 | 0 | 0.0 | 0.0 |
| capture-1 | sediment-61 | Positive | 38.36 | 6825 | 2150 | 2150 | 31.5 | 100.0 |
| capture-2 | sediment-61 | Positive | 38.36 | 9139 | 2801 | 2801 | 30.6 | 100.0 |
| capture-1 | sediment-62 | Positive | 32.11 | 290124 | 187124 | 156618 | 54.0 | 83.7 |
| capture-2 | sediment-62 | Positive | 32.11 | 304333 | 208477 | 152573 | 50.1 | 73.2 |
| capture-1 | sediment-63 | Positive | 32.18 | 202819 | 124051 | 83566 | 41.2 | 67.4 |
| capture-2 | sediment-63 | Positive | 32.18 | 196436 | 126576 | 88699 | 45.2 | 70.1 |
| capture-1 | sediment-64 | Positive | 35.26 | 1325 | 0 | 0 | 0.0 | 0.0 |
| capture-2 | sediment-64 | Positive | 35.26 | 1700 | 0 | 0 | 0.0 | 0.0 |
| capture-1 | sediment-65 | Positive | 35.40 | 50031 | 25932 | 18793 | 37.6 | 72.5 |
| capture-2 | sediment-65 | Positive | 35.40 | 43528 | 17321 | 12025 | 27.6 | 69.4 |
| capture-1 | sediment-66 | Positive | 36.38 | 4304 | 0 | 0 | 0.0 | 0.0 |
| capture-2 | sediment-66 | Positive | 36.38 | 5579 | 0 | 0 | 0.0 | 0.0 |
| capture-1 | sediment-67 | Positive | 38.45 | 70801 | 46010 | 18791 | 26.5 | 40.8 |
| capture-2 | sediment-67 | Positive | 38.45 | 78229 | 52760 | 16894 | 21.6 | 32.0 |
| capture-1 | sediment-68 | Positive | 38.62 | 31988 | 13962 | 8058 | 25.2 | 57.7 |
| capture-2 | sediment-68 | Positive | 38.62 | 28440 | 10387 | 3771 | 13.3 | 36.3 |
| capture-1 | sediment-69 | Positive | 34.14 | 327166 | 198942 | 133144 | 40.7 | 66.9 |
| capture-2 | sediment-69 | Positive | 34.14 | 366586 | 233981 | 151472 | 41.3 | 64.7 |
| capture-1 | sediment-70 | Positive | 36.99 | 3377 | 0 | 0 | 0.0 | 0.0 |
| capture-2 | sediment-70 | Positive | 36.99 | 4653 | 0 | 0 | 0.0 | 0.0 |
| capture-1 | sediment-71 | Positive | 33.89 | 132503 | 86232 | 66531 | 50.2 | 77.2 |
| capture-2 | sediment-71 | Positive | 33.89 | 136011 | 87949 | 74312 | 54.6 | 84.5 |
| capture-1 | sediment-72 | Positive | 36.30 | 84741 | 57300 | 57300 | 67.6 | 100.0 |
| capture-2 | sediment-72 | Positive | 36.30 | 84098 | 54494 | 36013 | 42.8 | 66.1 |
| capture-1 | sediment-73 | Positive | 37.02 | 32700 | 20535 | 4581 | 14.0 | 22.3 |
| capture-2 | sediment-73 | Positive | 37.02 | 32030 | 19401 | 7514 | 23.5 | 38.7 |
| capture-1 | sediment-74 | Positive | 36.36 | 3011 | 0 | 0 | 0.0 | 0.0 |
| capture-2 | sediment-74 | Positive | 36.36 | 3948 | 0 | 0 | 0.0 | 0.0 |
| capture-1 | sediment-75 | Suspect |  | 42 | 0 | 0 | 0.0 | 0.0 |
| capture-2 | sediment-75 | Suspect |  | 30 | 0 | 0 | 0.0 | 0.0 |
| capture-1 | sediment-76 | Suspect |  | 110 | 0 | 0 | 0.0 | 0.0 |
| capture-2 | sediment-76 | Suspect |  | 83 | 0 | 0 | 0.0 | 0.0 |
| capture-1 | sediment-77 | Suspect |  | 5154 | 0 | 0 | 0.0 | 0.0 |
| capture-2 | sediment-77 | Suspect |  | 6996 | 0 | 0 | 0.0 | 0.0 |
| capture-1 | sediment-78 | Suspect |  | 4327 | 0 | 0 | 0.0 | 0.0 |
| capture-2 | sediment-78 | Suspect |  | 6691 | 681 | 0 | 0.0 | 0.0 |
| capture-1 | sediment-79 | Suspect |  | 22 | 0 | 0 | 0.0 | 0.0 |
| capture-2 | sediment-79 | Suspect |  | 23 | 0 | 0 | 0.0 | 0.0 |
| capture-1 | sediment-80 | Suspect |  | 24 | 0 | 0 | 0.0 | 0.0 |
| capture-2 | sediment-80 | Suspect |  | 11 | 0 | 0 | 0.0 | 0.0 |
| capture-1 | sediment-81 | Suspect |  | 8203 | 0 | 0 | 0.0 | 0.0 |
| capture-2 | sediment-81 | Suspect |  | 11354 | 0 | 0 | 0.0 | 0.0 |
| capture-1 | sediment-82 | Suspect | 41.08 | 3450 | 248 | 0 | 0.0 | 0.0 |
| capture-2 | sediment-82 | Suspect | 41.08 | 5799 | 1158 | 295 | 5.1 | 25.5 |
| capture-1 | sediment-83 | Suspect |  | 9056 | 0 | 0 | 0.0 | 0.0 |
| capture-2 | sediment-83 | Suspect |  | 12411 | 0 | 0 | 0.0 | 0.0 |
| capture-1 | sediment-84 | Suspect | 40.83 | 6461 | 705 | 705 | 10.9 | 100.0 |
| capture-2 | sediment-84 | Suspect | 40.83 | 7461 | 634 | 634 | 8.5 | 100.0 |
| capture-1 | sediment-85 | Suspect |  | 12275 | 4537 | 4537 | 37.0 | 100.0 |
| capture-2 | sediment-85 | Suspect |  | 15046 | 5034 | 5034 | 33.5 | 100.0 |
| capture-1 | sediment-86 | Suspect |  | 19177 | 7919 | 2106 | 11.0 | 26.6 |
| capture-2 | sediment-86 | Suspect |  | 17940 | 5936 | 661 | 3.7 | 11.1 |
| capture-1 | sediment-87 | Suspect |  | 3038 | 0 | 0 | 0.0 | 0.0 |
| capture-2 | sediment-87 | Suspect |  | 4335 | 0 | 0 | 0.0 | 0.0 |
| capture-1 | sediment-88 | Suspect |  | 22078 | 8490 | 8490 | 38.5 | 100.0 |
| capture-2 | sediment-88 | Suspect |  | 20958 | 4620 | 4620 | 22.0 | 100.0 |
| capture-1 | sediment-89 | Suspect |  | 18321 | 7915 | 4114 | 22.5 | 52.0 |
| capture-2 | sediment-89 | Suspect |  | 22887 | 8962 | 3379 | 14.8 | 37.7 |
| capture-1 | sediment-90 | Suspect |  | 8826 | 201 | 201 | 2.3 | 100.0 |
| capture-2 | sediment-90 | Suspect |  | 12281 | 403 | 403 | 3.3 | 100.0 |
| capture-1 | Undetermined |  |  | 987909 | 30901 | 0 | 0.0 | 0.0 |
| capture-2 | Undetermined |  |  | 989527 | 40473 | 0 | 0.0 | 0.0 |
| capture-1 | control-1 |  |  | 1727947 | 25754 | 0 | 0.0 | 0.0 |
| capture-2 | control-1 |  |  | 1697915 | 21957 | 0 | 0.0 | 0.0 |
| capture-1 | control-2 |  |  | 1251991 | 21795 | 0 | 0.0 | 0.0 |
| capture-2 | control-2 |  |  | 1219539 | 9900 | 0 | 0.0 | 0.0 |
| capture-1 | control-3 |  |  | 862722 | 2003 | 0 | 0.0 | 0.0 |
| capture-2 | control-3 |  |  | 877188 | 2514 | 0 | 0.0 | 0.0 |
| capture-1 | control-4 |  |  | 2156783 | 4287 | 0 | 0.0 | 0.0 |
| capture-2 | control-4 |  |  | 2115832 | 6104 | 0 | 0.0 | 0.0 |
| capture-1 | control-5 |  |  | 1277530 | 18481 | 0 | 0.0 | 0.0 |
| capture-2 | control-5 |  |  | 1302672 | 19157 | 0 | 0.0 | 0.0 |
| capture-1 | control-6 |  |  | 1596828 | 1234 | 0 | 0.0 | 0.0 |
| capture-2 | control-6 |  |  | 1577998 | 0 | 0 | 0.0 | 0.0 |

